# A therapeutically targetable NOTCH1-SIRT1-KAT7 axis in T-cell Leukemia

**DOI:** 10.1101/2022.05.21.492944

**Authors:** Olga Lancho, Amartya Singh, Victoria da Silva-Diz, Luca Tottone, Patricia Renck Nunes, Maya Aleksandrova, Jesminara Khatun, Shirley Luo, Caifeng Zhao, Haiyan Zheng, Eric Chiles, Zhenyu Zuo, Pedro P. Rocha, Xiaoyang Su, Hossein Khiabanian, Daniel Herranz

## Abstract

T-cell Acute Lymphoblastic Leukemia (T-ALL) is a NOTCH1-driven disease in need of novel therapies. Here, we identify a NOTCH1-SIRT1-KAT7 link as a therapeutic vulnerability in T-ALL, in which SIRT1 is overexpressed downstream of a novel NOTCH1-bound enhancer. SIRT1 loss impairs leukemia generation, while SIRT1 overexpression accelerates leukemia and confers resistance to NOTCH1 inhibition in a deacetylase-dependent manner. Moreover, secondary SIRT1 loss extends survival and synergizes with NOTCH1 inhibition. Global acetyl-proteomics upon SIRT1 loss uncovered hyperacetylation of KAT7 and BRD1, subunits of a histone acetyltransferase complex targeting H4K12. Metabolic and gene expression profiling revealed a metabolic crisis together with a transcriptional signature resembling KAT7 deletion. Consistently, SIRT1 loss resulted in reduced H4K12ac, and overexpression of a non-acetylatable KAT7 mutant partly rescued SIRT1 loss-induced proliferation defects. The newly unveiled NOTCH1-SIRT1-KAT7 axis uncovers novel therapeutic targets in T-ALL and reveals a circular feedback mechanism balancing deacetylase/acetyltransferase activation with potentially broad relevance in cancer.

**Statement of significance:** We identified a novel axis in T-ALL whereby NOTCH1 activates *SIRT1* through an enhancer region, and SIRT1 deacetylates and activates KAT7. Targeting SIRT1 shows antileukemic effects, partly mediated by KAT7 inactivation. Our results identify novel therapeutic targets and uncover a rheostat mechanism between deacetylase/acetyltransferase activities with potentially broader cancer relevance.

## Introduction

T-cell acute lymphoblastic leukemia (T-ALL) is a rare and aggressive hematological cancer characterized for its frequent infiltration of the central nervous system and other organs (1). T-ALL affects both children and adults, accounting for 10-15% of pediatric and 20-25% of adult ALL diagnosed cases (1). Patient outcomes in the last decades have vastly improved thanks to intensive chemotherapy regimens (2–4). Despite the refinement of these therapies, 20% of pediatric and 50% of adult T-ALL cases still experience drug resistance and eventually relapse, showing dismal clinical outcomes with cure rates less than 30%. In addition, patients who get cured typically suffer from long-term debilitating comorbidities (2,5–8), overall underscoring the need to identify novel therapeutic approaches for the treatment of this disease.

The discovery of highly prevalent activating mutations in NOTCH1 in 60% of the patients underscores the role of NOTCH1 as the main oncogenic driver in T-ALL (9). Hence, the discovery of γ-secretase inhibitors (GSIs), which effectively block NOTCH1 activation via inhibition of a critical intramembrane proteolytic cleavage required for NOTCH1 signaling (10), held promise for the treatment of relapsed and refractory T-ALL. However, the clinical development of anti-NOTCH1 therapies in T-ALL has been hindered by limited therapeutic response to these drugs in clinical trials, exasperated by on-target gut toxicity (11). In this setting, cancer-specific metabolic rewiring and epigenetic remodeling represent critical hallmarks of cancer (12–14), and previous studies from our lab and others have specifically demonstrated the importance of both metabolic (15) and epigenetic (16) mechanisms mediating resistance to NOTCH1 inhibition *in vivo*. Thus, we postulated that central regulators that control both the metabolic and epigenetic status of cells could act as master regulators of NOTCH1-induced transformation and yield novel therapeutic targets in T-ALL.

In this context, we investigated the role of the NAD^+^-dependent SIRT1 histone deacetylase in T-ALL. SIRT1 is a well-known metabolic and epigenetic master regulator (17–23) which deacetylates H4K16ac and H3K9ac (20,24,25) and also regulates glycolysis and lipolysis through its complex interactions with PGC1α (26), HIF2 α (27), PPARγ (28) and AMPK (29, 30). In addition, SIRT1 has been shown to deacetylate key proteins in T-ALL, such as NOTCH1 (31), MYC (32), AKT (33) or PTEN (34), and it has been linked to deacetylation of other relevant proteins in cancer such as p53 (35, 36) or CDK2 (37). Specifically related to T-cells, previous literature showed that SIRT1 loss in the hematopoietic compartment failed to affect normal T-cell development (38). Regarding its role in hematological malignancies, SIRT1 has been previously shown to play both oncogenic or tumor suppressor roles in different contexts depending on disease-type and oncogenic driver. Indeed, SIRT1 was shown to promote leukemogenesis in Chronic Myeloid Leukemia (CML) (39, 40) and FLT3-ITD Acute Myeloid Leukemia (AML) (41). On the other hand, SIRT1 activation *in vivo* showed antiproliferative effects against MLL-rearranged leukemia (42) and myelodysplastic syndrome (MDS) (43). However, the potential role of SIRT1 in T-ALL *in vivo* is not well understood.

Here, we dissected the role of SIRT1 in T-ALL using a comprehensive approach that combined the use of primary mouse models with different genetic dosages of *Sirt1* with transcriptional and epigenetic profiling analyses from both human and mouse leukemias, as well as metabolomic and acetyl-proteomic analyses upon SIRT1 loss in T-ALL *in vivo*.

## Materials and methods

### Cell lines and culture conditions

Cells were cultured in standard conditions in a humidified atmosphere in 5 % CO_2_ at 37C in HyClone RPMI 1640 Media (Fisher Scientific, SH3002701) with 10 % FBS (Gemini Bio-Products, 900-108) and 100 U/mL penicillin and 100 μg/mL streptomycin (VWR, 45000-652). DND41 (ACC 525), HPB-ALL (ACC 483) and JURKAT (ACC 282) cells were obtained from Deutsche Sammlung von Mikroorganismen und Zellkulturen (DSMZ). Tumor-derived cell lines from mouse primary T-ALLs were generated and cultured in Opti-MEM (Life Technologies, 57985091) with 10% FBS, as previously described (44). Drugs used for different experiments include EX-527 (Sigma, E7034), DBZ (Syncom, 29762), WM-3835 (Tocris, 7366), 4-Hydroxytamoxifen (Sigma, H7904), doxycycline (Sigma, D9891), puromycin (Sigma, P8833), methyl-pyruvate (Sigma, 371173) and dimethyl-2-oxoglutarate (Sigma, 349631).

### Cell survival experiments

To analyze cell survival, cells were plated in triplicates on 24 well plates. Cell lines were seeded at 300,000 cells per well in a final volume of 1 mL RPMI 1640 media under the experimental conditions described in each experiment. 1 μM 4-Hydroxitamoxifen was added to induce *Sirt1* deletion, when indicated. 1 μg/ml doxycycline was added to induce *SIRT1* knockdown, when indicated. 250 nM DBZ was used to inhibit NOTCH1, when indicated. 90 μM EX-527 was used to inhibit SIRT1, when indicated. Every 3 days of treatment, cells were counted using a Countess II FL instrument (Fisher Scientific).

### Cell viability assays and evaluation of synergisms

Viability and cell growth ratios were determined analyzing cell density by MTT using the Cell Proliferation Kit I (Roche, 11465007001) in DND41 cells treated with EX-527 (10^-6^ to 10^-3^ M) for 3 and 6 days. Synergism was evaluated using the Chou-Talalay method in DND41 cells treated with EX-527, DBZ or with the combination of both EX-527 and DBZ using doses equivalent to 0.25 x IC50; 0.5 x IC50; 1 x IC50; 2 x IC50 or 4 x IC50 to calculate synergy (EX-527 IC50= 90 μM; DBZ IC50= 250 nM). Isobolograms were used to graphically represent the interaction between the two drugs and the Combination Index (CI) was determined using the Calcusyn software package (Biosoft) (45).

### Human cell transductions

Doxycycline-inducible GFP-expressing lentiviral vectors either non-targeting (shControl, VSC11521) or targeting *SIRT1* (V3SH11252-225149601 or V3SH11252_230472600) were obtained from Dharmacon. DND41 cells were transduced with each construct and selected with 5 μg/ml puromycin. Selected cells were treated with 1 μg/ml doxycycline to induce the different shRNAs together with concomitant GFP expression for downstream analyses.

For KAT7 overexpression experiments, we first obtained the *Kat7* cDNA from Genecopoiea (EX-Mm30842-Lv224-GS). K277Q or K277R directed mutagenesis was performed on this vector using the QuikChange II XL Site-Directed Mutagenesis Kit (Agilent, 200521), following manufacturer’s instructions, and constructs were sequence verified. Then, we subcloned the different *Kat7* constructs in the multiple cloning site of the pMSCV-mCherry FP vector (Addgene, 52114). Finally, empty vector or different *Kat7*- containing vectors were used to transduce and sort the tumor-derived *Sirt1* conditional knockout leukemia cell line using an Influx High Speed Sorter (BD Biosciences). Sorted cells were then used for downstream analyses.

### CRISPR/Cas9-induced loss of N-Se

Custom Alt-R CRISPR-Cas9 crRNA targeting sequences both flanks of the N-Se region (5’-GTTCATCGGTGGAGTTGAGG-3’ and 5’-AGCGTTTTCAAGTTATAATC-3’, respectively) were obtained from Integrated DNA Technologies (IDT) and used to generate JURKAT N-Se^-/-^ cells following the manufacturer’s instructions. Briefly, each crRNA and Alt-R CRISPR-Cas9 tracrRNA-ATTO 550 (IDT, 1075927) oligos were mixed in equimolar concentrations to a final duplex concentration of 44 μM. Annealing was achieved by heating up the mixture to 95 °C for 5 min and subsequent slow cooling down to room temperature (20-25C). 22 pmol of each crRNA:tracrRNA duplex were incubated with 18 pmol of Alt-R S.p. Cas9 Nuclease V3 (IDT, 1081059) for 20 minutes at room temperature. After the formation of the crRNA:tracrRNA:Cas9 complexes, the complexes targeting the 5’ and 3’ end of N-Se were mixed in equimolar concentrations. 0.5 million JURKAT cells were electroporated with 40 pmol of crRNA:tracrRNA:Cas9 complexes using the Neon Transfection System with 10 μL electroporating tips (MPK1096, Thermo Fisher Scientific) in duplicate. Electroporation conditions were 3 pulses at 1600 V and 10 ms width. ATTO 550-positive single-cell clones were sorted using the Influx High Speed Sorter (BD Biosciences) 24 h after transfection. Clones were screened for N-Se loss by PCR using REDTaq ReadyMix (Sigma, R2523-100RXN) and custom primers (5’-AGCGAGACTCCGTCTAGAAA-3’, 5’-CTCGAACTCCTGACCTCAATC-3’ and 5’-CAGCAAAATTGAGGGGAAAG-3’).

### CRISPRa and CRISPRi experiments

pLV-U6-gRNA-UbC-DsRed-P2A-Bsr plasmid (Addgene, 83919) was used to express gRNAs targeting human *SIRT1* TSS and N-Se. Assembly of custom lentiviral vectors expressing sgRNAs of choice was accomplished by Golden Gate cloning and type IIS restriction enzyme BsmBI (NEB, R0739), which cleaves outside its recognition sequence to create unique overhangs as previously reported (46). Primers used to clone *SIRT1* TSS gRNA: sense primer 5′-CACCGGGGCAGCCAAATTCGCCCCT-3′ and antisense primer 5′-AAACAGGGGCGAATTTGGCTGCCCC-3′. Primers used to clone N-Se gRNA1: sense primer 5′-CACCGGGCTCCAGAAGCTCCGAGCG-3′ and antisense primer 5′-AAACCGCTCGGAGCTTCTGGAGCCC-3′. Primers used to clone N-Se gRNA2: sense primer 5′-CACCGGGGCCTCCTTTGCTCGTATT-3′ and antisense primer 5′-AAACAATACGAGCAAAGGAGGCCCC-3′.

pLV hUbC-dCas9 VP64-T2A-GFP lentiviral vector expressing dCas9VP64 (Addgene, 53192) or pLV hUbC-dCas9 KRAB-T2A-GFP lentiviral vector expressing dCas9KRAB (Addgene, 67620) were used to transduce DND41 cells, followed by selection of GFP- positive cells by fluorescence-activated cell sorting (FACS) using an Influx High Speed Sorter (BD Biosciences), in order to have DND41 stably expressing either dCas9-VP64-GFP or dCas9-KRAB-GFP, respectively. Subsequently, one lentiviral vector encoding for each gRNA-DsRed was used to transduce DND41-dCas9VP64-GFP+ or DND41-dCas9KRAB-GFP+ models. Cells co-expressing both GFP and DsRed markers (>90%) were further isolated by FACS and processed for the evaluation of SIRT1 expression.

### Luciferase Reporter Assays

We performed reporter assays using a pGL4.10 Vector (Promega, E6651) luciferase construct alone or coupled with the N-Se enhancer sequence (hg19; chr10:69,608,815–69,610,235; DNA sequence was synthesized by Genewiz), cloned in the forward and reverse orientations, and following our previously described protocol (47). Briefly, we electroporated JURKAT cells with a Neon Transfection System Device (MPK5000, Thermo Fisher Scientific) using 100 μL tips (MPK10025, Thermo Fisher Scientific). Electroporation conditions were as follows: pulse voltage 1,350 V, pulse width 10 ms, pulse number 3, and cell density 10^7^/mL. Constructs were transfected together with a plasmid driving the expression of the Renilla luciferase gene (pCMV-Renilla) used as an internal control. We measured luciferase activity 42 hours after electroporation with the Dual-Luciferase Reporter Assay kit (Promega, E1980).

### Oxygen Consumption Rate (OCR)

OCR rates were measured using a XF24 Seahorse Biosciences extracellular flux analyzer (Agilent Technologies) according to manufacturer’s instructions. Briefly, T-ALL cells were resuspended in Seahorse XF RPMI Medium (Agilent Technologies, 103576-100) supplemented with 10 mM glucose (Agilent Technologies, 1003577-100), 1 mM pyruvate (Agilent Technologies, 1003578-100) and 2 mM glutamine (Agilent Technologies, 1003579-100). 5 x 10^5^ cells per well were plated in XF24 Seahorse Biosciences plates pre-coated with Cell-Tak (Corning, 354240) and spun down on the plate to ensure that cells were completely attached. OCR was analyzed by sequential injections of 1 μM oligomycin (Sigma, O4876), 1 μM FCCP (Sigma, C2920) and 0.5 μM rotenone (Sigma, R8875) and antimycin A (Sigma, A8674) in each well.

### In vivo models of NOTCH1-driven mouse T-ALLs

Animals were maintained in ventilated caging in specific pathogen-free facilities at New Brunswick RBHS Rutgers Campus. All animal housing, handling, and procedures involving mice were approved by Rutgers Institutional Animal Care and Use Committee (IACUC), in accordance with all relevant ethical regulations.

To generate NOTCH1-induced *Sirt1*-overexpresing leukemias, we performed a retroviral transduction of lineage-negative enriched cells from *Sirt1*^WT^ or *Sirt1*^TG^ (48) (JAX, #024510) donors with retrovirus encoding ΔE-NOTCH1-GFP, an oncogenic activated form of NOTCH1 with concomitant expression of GFP, as previously described (15). Cells were then transplanted via retro-orbital injection into lethally irradiated (7.5 Gy) recipient mice. Investigators were not blinded to group allocation. Animals were monitored for signs of distress or motor function at least twice daily, until they were terminally ill, whereupon they were euthanized.

In order to generate *Sirt1* conditional inducible knockout (*Sirt1*^flox/flox^) leukemias, we first crossed *Sirt1*^flox/flox^ mice (38) (JAX, #029603) with mice harboring a tamoxifen-inducible Cre recombinase from the ubiquitous Rosa26 locus (49). Then, we performed a retroviral transduction of lineage-negative enriched cells from *Sirt1*^flox/flox^-Rosa26^Cre-ERT2/+^ donors with retrovirus encoding ΔE-NOTCH1-GFP or HDΔP-NOTCH1-GFP oncogenic activated forms of NOTCH1 with concomitant expression of GFP, as previously described (15). Cells were then transplanted via retro-orbital injection into lethally irradiated (7.5 Gy) recipient mice.

For leukemia-initiation survival studies, mice transplanted with ΔE-NOTCH1-GFP-infected *Sirt1*^flox/flox^-Rosa26^Cre-ERT2/+^ progenitors were treated 48 hours after transplantation with vehicle only (corn oil; Sigma, C8267) or tamoxifen (Sigma, T5648; 3 mg per mouse in corn oil), to induce isogenic loss of *Sirt1* before leukemic transformation.

For leukemia-progression studies, already generated E-NOTCH1-GFP-induced or HD P-NOTCH1-GFP-induced *Sirt1*^flox/flox^-Rosa26^Cre-ERT2/+^ leukemias (1 x 10^6^ leukemia cells) were transplanted from primary recipients into sub-lethally irradiated (4.5 Gy) 6-8- week-old secondary recipient C57BL/6 mice (Taconic Farms) by retro-orbital injection. 48 hours after leukemic cell transplantation, recipient mice were treated with vehicle only (corn oil; Sigma, C8267) or tamoxifen (Sigma, T5648; 3 mg per mouse in corn oil), to induce isogenic loss of *Sirt1* in established leukemias. Subsequently, 5 days post-transplantation, mice in each arm were divided randomly into two different groups: control groups were subsequently treated with vehicle only (2.3 % DMSO in 0.005% methylcellulose, 0.1 % Tween-80) or with DBZ (5 mg per Kg in vehicle solution) on a 4- day-ON and 3-day-OFF schedule, as previously described (15). Investigators were not blinded to group allocation. Animals were monitored for signs of distress or motor function at least twice daily, until they were terminally ill, whereupon they were euthanized.

To generate *Sirt1/Ampk double* conditional inducible knockout (*Sirt1*^flox/flox^; *Ampk*^flox/flox^-Rosa26^Cre-ERT2/+^) leukemias, we first crossed these mice with *Ampk*^flox/flox^ mice (50) (JAX, #014141), and performed leukemia-progression studies upon transplantation of already generated E-NOTCH1-GFP-induced *Sirt1*^flox/flox^;*Ampk*^flox/flox^-Rosa26^Cre-ERT2/+^ leukemias (1 x 10^6^ leukemia cells) similar to before.

For *Sirt1* acute deletion analyses in mouse primary leukemias (metabolomics, acetyl-proteomics, RNAseq, ChIPseq, western blot and/or apoptosis studies), we transplanted lymphoblasts from spleens of *Sirt1* conditional knockout NOTCH1-induced T-ALL-bearing mice into a secondary cohort of recipient mice, as before. We monitored mice until they presented clear leukemic signs with >60% GFP-positive leukemic cells in peripheral blood; then, mice were treated with vehicle or tamoxifen and, 48 hours after treatment, mice were euthanized and spleen samples were collected for further analyses.

For analysis of the effects of *Sirt1* overexpression on the response to DBZ *in vivo*, we infected NOTCH1-induced GFP-positive mouse primary leukemic cells with a retrovirus driving bicistronic expression of the cherry fluorescent protein together with either *Sirt1* (pMSCV-Sirt1-IRES-mCherry FP) or *Sirt1* H355A (pMSCV-Sirt1-H355A-IRES-mCherry FP). *Sirt1* cDNA was amplified by PCR, using primers containing unique restriction enzyme sites, cloned into the multiple cloning site of the pMSCV-mCherry FP vector (Addgene, 52114), and sequence verified. *Sirt1* H355A mutation was generated by performing directed mutagenesis on this vector using the QuikChange II XL Site-Directed Mutagenesis Kit (Agilent, 200521), following manufacturer’s instructions. A mix of transduced and non-transduced cells (i.e., without sorting) were then injected into sublethally irradiated C57BL/6 mice (4.5 Gy). Mice were treated with DBZ (5 mg per Kg) on daily basis and leukemic blasts in peripheral blood were analyzed as previously described (15).

### Western blotting

Human peripheral blood mononuclear cells (PBMCs) and peripheral blood CD4-positive cells were purchased from Lonza (product numbers CC-2702 and 2W-200, respectively). Samples from normal human thymus were kindly provided by Dr. Adolfo Ferrando (Columbia University). Whole-cell extracts were prepared using standard procedures. After protein transfer, membranes were incubated with the antibodies anti-SIRT1(1:1000, sc-74504, Santa Cruz), anti-Sirt1 (1:5000, ab12193, Abcam), anti-AMPK (1:1000, 5831S, Cell Signaling), anti-p-AMPK (1:1000, 2535S, Cell Signaling), anti-4E-BP1 (1:1000, 9644S, Cell Signaling), anti-p-4EB-P1 (1:1000, 2855S, Cell Signaling), anti-p53 (1:1000, 2524S, Cell Signaling), anti-ac-p53 (1:1000, 2570S, Cell Signaling), anti-ac-NFkB (1:1000, ab191870, Abcam), anti-H4 (1:1000, ab10158, Abcam), anti-H4K12ac (1:1000, ab177793, Abcam), anti-GAPDH (1:10000, 97166, Cell Signaling) and anti-β-actin-HRP (1:50000; A3854, Sigma). Antibody binding was detected with a secondary antibody coupled to horseradish peroxidase (Sigma, NA934 and NA931) using enhanced chemiluminescence (Thermo Scientific, 34578) or using secondary fluorescence-coupled antibodies (Thermo Scientific) in an iBright FL1500 instrument (Thermo Scientific). Immunoblot quantification was performed using Fiji image processing package (51).

### Flow cytometry analysis

To analyze spleen samples, single cell suspensions were prepared by disrupting spleens through a 70 μm filter. Red cells were removed by incubation with ammonium-chloride-potassium lysing buffer (155 mM NH4Cl, 12 mM KHCO3 and 0.1 mM EDTA) for 5 minutes in ice. Apoptotic cells in leukemic spleens or human/mouse cell lines after different treatments were quantified with PE-AnnexinV Apoptosis Detection Kit I (BD Pharmingen, 559763). All flow cytometry data was collected on an Attune NxT Flow Cytometer (ThermoFisher Scientific) and analyzed with FlowJo v10.6.2 software (BD).

### Metabolite extraction

For leukemic spleens, leukemic mice were euthanized, and spleens were quickly removed, minced and frozen. Subsequently, 25 mg per sample of frozen leukemic spleens were homogenized under liquid nitrogen flow using a Retsch CryoMill at 20 Hz for 2 min. Pulverized samples were then mixed with methanol:acetonitrile:water (40:40:20) with 0.1 M formic acid solution followed by 10 min incubation on ice and 500 μL of the extract was neutralized with 44 μL of 15 % (m/v) ammonium bicarbonate. Finally, after centrifugation (14,000 g, 10 min at 4 °C), samples were transferred to clean tubes and sent for LC-MS analysis. For media, we diluted 20 μL of each respective media into 980 μL of ice-cold solvent (40:40:20 methanol:acetonitrile:water + 0.1 M formic acid) followed by neutralization with 80 μL of 15 % ammonium bicarbonate. Samples were directly submitted to LC-MS analysis.

### LC-MS-based metabolomics

LC−MS analysis of the extracted metabolites was performed on a Q Exactive PLUS hybrid quadrupole-orbitrap mass spectrometer (ThermoFisher Scientific) coupled to hydrophilic interaction chromatography (HILIC). The LC separation was performed on Vanquish Horizon UHPLC system with an XBridge BEH Amide column (150 mm x 2.1 mm, 2.5 μM particle size, Waters, Milford, MA) with the corresponding XP VanGuard Cartridge. The liquid chromatography used a gradient of solvent A (95 %:5 % H2O:acetonitrile with 20 mM ammonium acetate, 20 mM ammonium hydroxide, pH 9.4), and solvent B (20%:80% H2O:acetonitrile with 20 mM ammonium acetate, 20 mM ammonium hydroxide, pH 9.4). The gradient was 0 min, 100% B; 3 min, 100% B; 3.2 min, 90% B; 6.2 min, 90% B; 6.5 min, 80% B; 10.5 min, 80% B; 10.7 min, 70% B; 13.5 min, 70% B; 13.7 min, 45% B; 16 min, 45% B; 16.5 min, 100% B. The flow rate was 300 μl/min. Injection volume was 5 μl and column temperature 25 °C. The MS scans were in negative ion mode with a resolution of 70,000 at m/z 200. The automatic gain control (AGC) target was 3 x 10^6^, and the scan range was 75-1,000. Metabolite features were extracted in MAVEN (52) with the labeled isotope specified and a mass accuracy window of 5 ppm.

### Acetyl Proteomics sample preparation

Leukemic mice (E-NOTCH1-GFP-induced or HD P-NOTCH1-GFP-induced *Sirt1*^flox/flox^-Rosa26^Cre-ERT2/+^ leukemia-bearing mice) were euthanized, and spleens were quickly removed, minced and frozen. Subsequently, 25 mg per sample of frozen leukemic spleens were homogenized under liquid nitrogen flow using Retsch CryoMill at 20 Hz for 2 min. Pulverized samples were then mixed with 2x laemmli buffer with 50 mM DTT in a ratio of 1 mg of tissue: 10 ml of 2x laemmli buffer. Samples were sonicated for 7 min, incubated at 95 °C for 10 min and centrifuged (25,000 g, 30 min at 4 °C), supernatant was saved in a new tube. Equal volume of 8 M Urea was used to extract the remaining precipitate with sonication and centrifugation (25,000 g, 30 min at 4 °C), supernatant was combined with the previous one and protein concentration was determined by 660 kit (ThermoFisher Scientific). 500 µg of lysate was run into SDS-PAGE gel and digested using standard gel-plug procedures. 450 µg of each sample were labeled with TMT10 plex reagent (ThermoFisher Scientific) according to manufacturer’s instructions. A test mix of 1% of individual samples were run on nano-LC-MSMS and the reporter ion intensity was used to normalize the mixing of the samples to achieve a ratio of 1:1 for all samples. After mixing, the samples were desalted with SpeC18 (Waters) and dried. TMT10 labeled Peptides with acetylated K were enriched with Acetyl-Lysine Motif kit (Cell Signaling Technology) following manufacturer’s instructions. The eluted peptides were desalted with Stagetips (53).

### Acetyl Proteomics Nano-LC-MS/MS

Nano-LC-MS/MS was performed using a Dionex rapid-separation liquid chromatography system interfaced with a QExactive HF (ThermoFisher Scientific). Samples were loaded onto an Acclaim PepMap 100 trap column (75 µm x 2 cm, ThermoFisher Scientific) and washed with Buffer A (0.1 % trifluoroacetic acid) for 5 min with a flow rate of 5 µl/min. The trap was brought in-line with the nano analytical column (nanoEase, MZ peptide BEH C18, 130A, 1.7 µm, 75 µm x 20 cm, Waters) with flow rate of 300 nL/min with a multistep gradient (4 % to 15 % buffer B [0.16 % formic acid and 80 % acetonitrile] for 20 min, then 15 % – 25 % B for 40 min, followed by 25 % – 50 % B for 30 min). Mass spectrometry data was acquired using a data-dependent acquisition procedure with a cyclic series of a full scan acquired with a resolution of 120,000 followed by tandem mass spectrometry scans (33 % normalized collision energy (NCE) 33% in the higher-energy collisional dissociation (HCD) cell with resolution of 45,000 of the 20 most intense ions with dynamic exclusion duration of 20 seconds).

Some samples were analyzed using a Dionex rapid-separation liquid chromatography system interfaced with a Orbitrap Eclipse tribrid mass spectrometer (ThermoFisher Scientific). The LC-conditions were the same as described for Dionex RSLC connectee to QExactive HF (ThermoFisher Scientific). The scan sequence began with an MS1 spectrum (Orbitrap analysis, resolution 120,000 with scan range from 375–1600 Th, automatic gain control (AGC) target 6E4, maximum injection time 50 ms). The top S (3 seconds) duty cycle scheme were used to determine the number of parent ions investigated for each cycle. For proteomic samples without IAP, SPS method was used. In detail, peptide ions were first collected for MS/MS by collision-induced dissociation (CID) and scanned in ion trap with AGC 2E4, NCE 35 %, maximum injection time 50 ms, and isolation window at 0.7. Following acquisition of each MS2 spectrum, 10 MS2 fragment ions were captured as the MS3 precursor population using isolation waveforms with multiple frequency notches. MS3 precursors were fragmented by higher-energy collisional dissociation (HCD) and analyzed using the Orbitrap (NCE 65, AGC 1.5E5, maximum injection time 105 ms, resolution 50,000 at 400 Th and scan range from 100-500 amu). For acetyl-peptide, MS/MS method was used. Parent mass was isolated in the quadrupole with an isolation window of 0.7 m/z, AGC target 1.75E5, and fragmented with HCD NCE38.The fragments were scanned in Orbitrap with resolution of 50,000. The MS/MS scan ranges were determined by the charge state of the parent ion but lower limit was set at 110 amu.

### Acetyl Proteomics Data analysis

LC-MS data was analyzed with Maxquant (version 1.6.17.0) with Andromeda search engine. For samples run on QExactive, Type of LC-MS run was set to reporter ion MS2 with 10plex TMT as isobaric labels. For data acquired on Eclipse, Type of LC-MS run was set to reporter ion MS3 with TMT10 plex as isobaric labels for proteome and MS2 with TMT10 plex for acetylome. Reporter ion mass tolerance was set at 0.003 Da. LC-MS data was searched against Uniprot mouse database with addition of potential contaminants. Protease was set as trypsin/P that allowed 2 miss cuts for total proteomic data and 5 miss cuts for acetyl K sample (PTM sample). Carbaidomethylation of cysteine was set as fixed modification, N-terminal acetylation, oxidation at methionine as well as acetylation at lysine were set as variable modifications. For quantification, spectra were filtered by min reporter PIF set at 0.5 (spectra purity) for Acetyl peptides and 0.75 for proteome data.

The Maxquant search results were analyzed using Perseus (version 1.6.15.0). Data was first filtered for reverse and contaminant hits and the reporter ion intensity data were further log 2 transformed. The proteome data was then normalized to median of each channel. For acetyl peptides, the data was also normalized based on proteome summed intensity of each channel and each acetyl site was further normalized to corresponding protein group quantitation value if the group had been identified in proteome data. For group comparison, statistical significance between groups was analyzed using student *t*-test with equal variance on both sides, requiring two valid values in at least one group, and the Q value was calculated with permutation. FDR was analyzed using student *t*-test with equal variance on both sides and the Q value was calculated using permutation test. Significance was assigned with consideration of S0 (S 0 = 0.1).

### Quantitative RT-PCR

Total RNA was extracted from cells using RNeasy Plus Mini Kit (Qiagen), and cDNA was generated with High-Capacity cDNA Reverse Transcription Kit (Applied Biosystems). Quantitative PCR was performed on a QuantStudio 3 Real-Time PCR System (Applied Biosystems) using SYBR Green PCR Master Mix (Takara), and transcript levels were normalized to actin as an internal control. Primers used: *SIRT1* (Forward: 5′-AGGATAGAGCCTCACATGCAA-3′; Reverse: 5′-TCGAGGATCTGTGCCAATCATA-3′), *Sirt1* (Forward: 5′-GAGCTGGGGTTTCTGTCTCC-3′; Reverse: 5′-CCGCAAGGCGAGCATAGATA-3′), *ACTIN* (5′-CAACGCCAACCGCGAGAAGAT-3′; Reverse: 5′-CCAGAGGCGTACAGGGATAGCAC-3′) and *Actin* (5′-GGCTGTATTCCCCTCCATCG-3′; 5′-CCAGTTGGTAACAATGCCATGT-3′).

### RNA-seq gene expression profiling

*Sirt1* conditional knockout E-NOTCH1-induced T-ALL-bearing mice were treated with vehicle only (corn oil; Sigma, C8267) or tamoxifen (Sigma, T5648; 3 mg per mouse in corn oil) to induce isogenic loss of *Sirt1* via intraperitoneal injection. 48 hours later, single-cell suspensions of total leukemic splenocytes were prepared by pressing leukemic spleens through a 70 μm filter. We removed red cells in spleen samples by incubation with red blood cell lysis buffer (155 mmol/L NH4Cl, 12 mmol/L KHCO3, and 0.1 mmol/L EDTA) for 5 minutes on ice. RNA was extracted using QIAshredder (QIAGEN, 79656) and RNeasy Mini (QIAGEN, 74106) kits. RNA library preparations and next-generation sequencing were performed using Illumina Next-Seq platform (Illumina). We estimated gene-level raw counts using kallisto 0.44.0 (54), with the Ensembl GRCm38 transcriptome as the reference. We evaluated differential expression between the samples in R using DESeq2 (55). GSEA analyses were performed to assess enrichment of KAT7-loss signature (56) based on 10,000 permutations of the gene list (57).

### ChIP-seq analysis

Analyses of genome-wide H4K12ac mark in leukemic cells isolated from *Sirt1* conditional knockout E-NOTCH1-induced T-ALL-bearing mice acutely treated as before with vehicle or tamoxifen to induce isogenic loss of *Sirt1* was done by Active Motif, following well-established protocols and using a ChIP-validated H4K12ac antibody (Active Motif, #39165). Peaks were called using either the MACS (58) or SICER algorithms (59). MACS default cutoff is *P* value 1e-7 for narrow peaks and 1e-1 for broad peaks, and SICER default cutoff is FDR 1e-10 with gap parameter of 600 bp. Peak filtering was performed by removing false ChIP-Seq peaks as defined within the ENCODE blacklist (60).

### RNA-seq analysis of Normal vs T-ALL

*SIRT1* expression was analyzed among T-ALL samples (n=57) and physiological thymocyte subsets (n=21) from published literature (61). Quantile normalization was performed across samples. Differential expression was performed using Mann-Whitney U-Test (P<0.001), and genes with FDR<0.05 using the Benjamini-Hochberg correction were shortlisted as differentially expressed.

### Statistical analysis

Statistical analyses were performed with Prism 8.0 (GaphPad). Unless otherwise indicated in figure legends, statistical significance between groups was calculated using an unpaired two-tailed Student’s *t*-test. Survival in mouse experiments was represented with Kaplan–Meier curves, and significance was estimated with the log-rank test.

### Data availability

RNA-seq and ChIP-seq data from *Sirt1* conditional knockout leukemias was deposited in the NCBI Gene Expression Omnibus repository under the following accession numbers: GSE203387 and GSE203386. We analyzed NOTCH1 transcriptional targets using GSI washout experiments publicly available from GEO: GSE29544. We analyzed *SIRT1* promoter occupancy of chromatin marks and epigenetic and transcription factors using the following T-ALL publicly available ChIP-seq and ATAC-seq datasets from Gene Expression Omnibus (GEO): GSE58406, GSE124223, GSE 29611, GSE54379, GSE29600 and GSE138516.

## Results

### Sirt1 upregulation in T-ALL is driven by a NOTCH1-bound enhancer

We first investigated the expression of *SIRT1* in ALL cell lines taking advantage of the Cancer Cell Line Encyclopedia (CCLE) (62). These analyses revealed that *SIRT1* levels in ALL are the highest among the different types of leukemias analyzed, including AML, CLL and CML samples (**Supplementary Fig. S1A**). Subsequent analyses of gene expression profiling data from pediatric T-ALL patients (61) revealed a significant upregulation of S*IRT1* expression in T-ALL compared to healthy thymocyte subsets (**Fig. 1A**). Interestingly, *SIRT1* levels were similar across different T-ALL clinical subgroups (63), with HOXA-driven T-ALLs showing relatively lower levels (**Supplementary Fig. S1B**). In line with this, we found SIRT1 protein levels to be significantly upregulated in T-ALL cells compared to healthy human peripheral mononuclear cells, CD4^+^ T-cells or human thymus samples (**Fig. 1B**). *SIRT1* is not known to be duplicated or mutated in T-ALL (63). Since NOTCH1 activating mutations are observed in ∼60% of T-ALL patients (9), these results led us to the hypothesize that NOTCH1 might be regulating *SIRT1* expression. Arguably, the most accurate way to uncover NOTCH1 direct targets is to use GSI washout methods (64). These assays allow NOTCH1 reactivation in the presence of cycloheximide (CHX) or dominant-negative MAML1 (DN-MAML1, a specific inhibitor of NOTCH1 signaling) upon removal of GSI (64). High-confidence direct NOTCH1 targets are defined by a ≥2-fold increase within 4h of GSI washout that is insensitive to CHX (independent of new protein translation) but sensitive to DN-MAML1 (dependent on NOTCH1 binding). Notably, *SIRT1* expression in T-ALL cells increased ∼4-fold after GSI removal, and this effect was insensitive to CHX but was rescued by DN-MAML1 (**Fig. 1C**), demonstrating that *SIRT1* is a direct target of NOTCH1 in T-ALL. Indeed, short-term treatment with GSIs in NOTCH1-driven human or mouse T-ALL cell lines *in vitro* led to a slight but consistent downregulation of SIRT1 protein levels (**Fig. 1D**). Epigenetic profiling analyses revealed that NOTCH1 binding to the *SIRT1* promoter in T-ALL was modest; however, it uncovered a discrete NOTCH1-binding peak located ∼35Kb upstream of the *SIRT1* promoter (**Fig. 1E**). Importantly, NOTCH1 binding to this region was shown in both CUTLL1 and HPB-ALL, two independent T-ALL cell lines. This region also exhibited additional features of NOTCH1 canonical signaling, such as the binding of RBPJ or ETS1, as well as bona-fide enhancer marks such as enrichment of H3K27ac and binding of BRD4 and ZNF143 (**Fig. 1D**). In addition, ATAC-seq data of two independent human primary T-ALL samples revealed that this region shows accessible chromatin (**Fig. 1E**). Thus, we postulated that this region (which we named N-Se, for **N**OTCH1-bound **S**IRT1 **e**nhancer) may act as an enhancer of *SIRT1* driven by NOTCH1, in an analogous way to our previously identified N-Me enhancer in the control of *MYC* (44).

**Figure 1.**
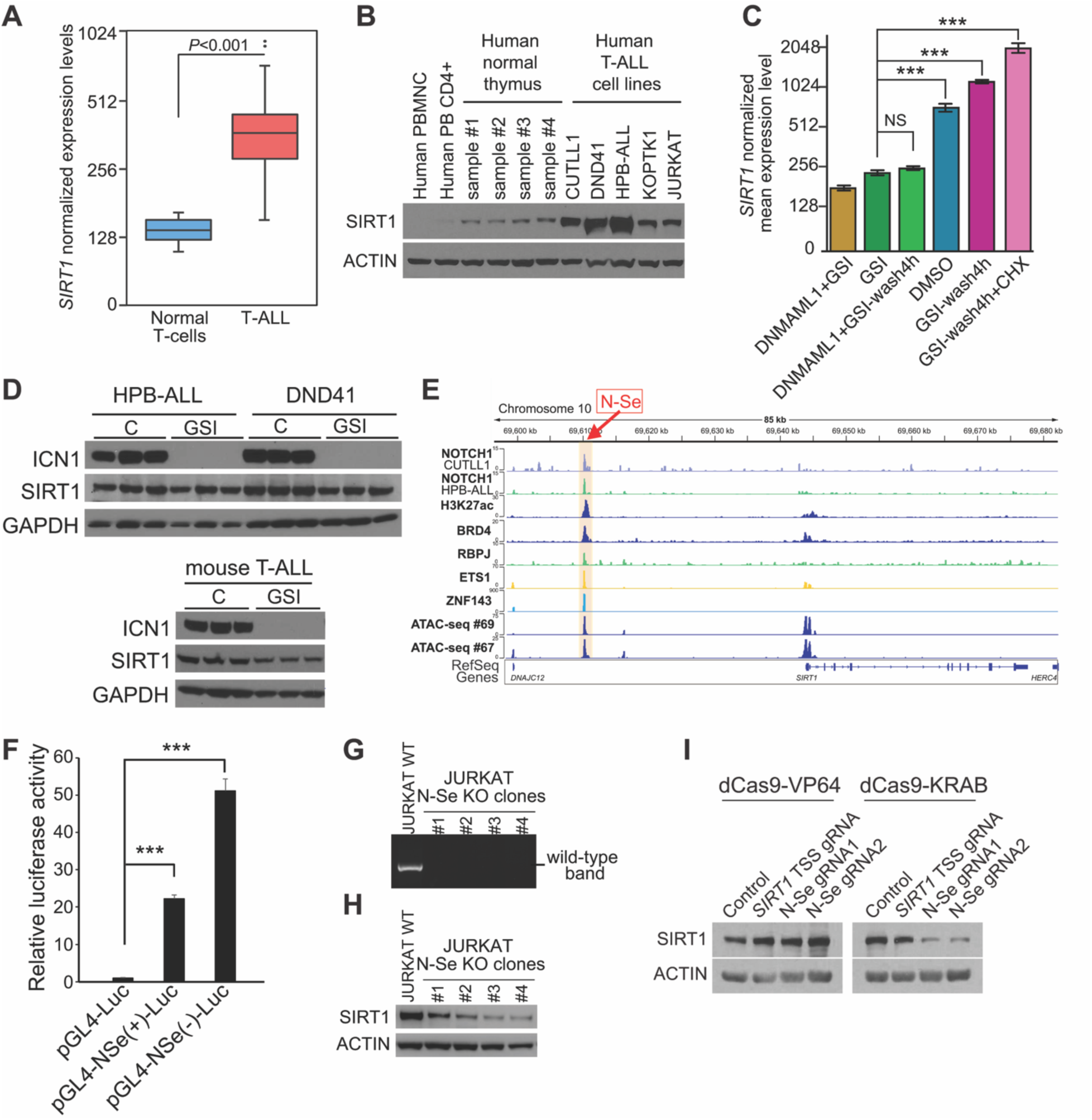
SIRT1 is overexpressed in T-ALL downstream of a NOTCH1-bound enhancer. **A,** Box-plot showing *SIRT1* expression among T-ALL samples (n=57) and physiological thymocyte subsets (n=21) (61). Quantile normalization was performed across samples. Boxes represent first and third quartiles and the line represents the median. Whiskers represent the upper and lower limits (*P*<0.001 using Mann-Whitney U-Test; FDR<0.05 using Benjamini-Hochberg correction). **B,** Western blot analysis of SIRT1 and ACTIN expression in human peripheral blood mononuclear cells (PBMNC), CD4+ T-cells or normal human thymocytes, as compared to human T-ALL cell lines. **C,** GSI washout experiments in CUTLL1 T-ALL cells, treated with GSI (Compound E, 1μM) for 3 days, washed twice, and incubated 4h in the presence or absence of 20μM cycloheximide (64). To control for GSI “off-NOTCH” effects, cells were also transduced with a dominant-negative MAML1 (DN-MAML1). (n=3 per condition; \*\*\**P* < 0.005 using two-tailed Student *t*-test; NS, not significant). **D,** Western blot analysis of NOTCH1 (ICN1), SIRT1 and ACTIN expression in triplicates from DND41 or HPB-ALL human T-ALL cells treated with DBZ (250nM) for 3 days or mouse T-ALL cells treated with DBZ (250nM) for 24h. **E,** Epigenetic profiling around the *SIRT1* promoter in human T-ALL showing ChIP-seq tracks in human T-ALL cell lines and ATAC-seq tracks in human T-ALL primary samples. N-Se enhancer highlighted in orange. **F,** Luciferase reporter activity in JURKAT cells of a pGL4 promoter empty construct (pGL4-Luc), a pGL4 promoter plus the human N-Se enhancer in the forward (NSe(+)-Luc) or reverse (NSe(-)-Luc) orientation. Data from three independent electroporation replicates are shown. \*\*\**P* < 0.005 using two-tailed Student *t*-test. **G,** Genotyping of JURKAT single-cell clones harboring a N-Se homozygous deletion. JURKAT cells not electroporated (WT) are shown as controls. **H,** SIRT1 protein expression levels via western blot analysis in JURKAT control cells or four independent JURKAT single-cell clones with N-Se homozygous deletion. **I,** SIRT1 protein expression levels via western blot analysis in DND41 cells harboring either a dCas9-VP64 or dCas9-KRAB construct, and infected with gRNAs targeting either the *SIRT1* promoter transcriptional start site (TSS) or two independent gRNAs targeting N-Se.

Luciferase reporter assays in T-ALL cells testing this hypothesis showed strong, orientation-independent activation of reporter constructs containing N-Se (**Fig. 1F**). Moreover, CRISPR/Cas9-induced deletion of N-Se resulted in reduced SIRT1 expression human T-ALL JURKAT cells (**Fig. 1G-H**). Finally, CRISPRi or CRISPRa assays targeting N-Se in DND41 T-ALL cells led to decreased or upregulated SIRT1 levels, respectively, to a similar extent as targeting the *SIRT1* promoter itself (**Fig. 1I**). These results demonstrate that this newly identified distal regulatory region controls *SIRT1* expression downstream of NOTCH1, which might underlie the broad upregulation of SIRT1 observed in T-ALL patients.

### Genetic or pharmacological inhibition of SIRT1 in vitro shows antileukemic effects

We next hypothesized that targeting SIRT1 might result in antileukemic effects in human T-ALL. Indeed, knockdown of *SIRT1* in DND41 T-ALL cells with two independent doxycycline-inducible shRNAs led to reduced SIRT1 levels **(Fig. 2A-B)**, together with a concomitant reduction in proliferation **(Fig. 2C)** and cytotoxic effects, as revealed by increased apoptosis **(Fig. 2D-E)**.

**Figure 2.**
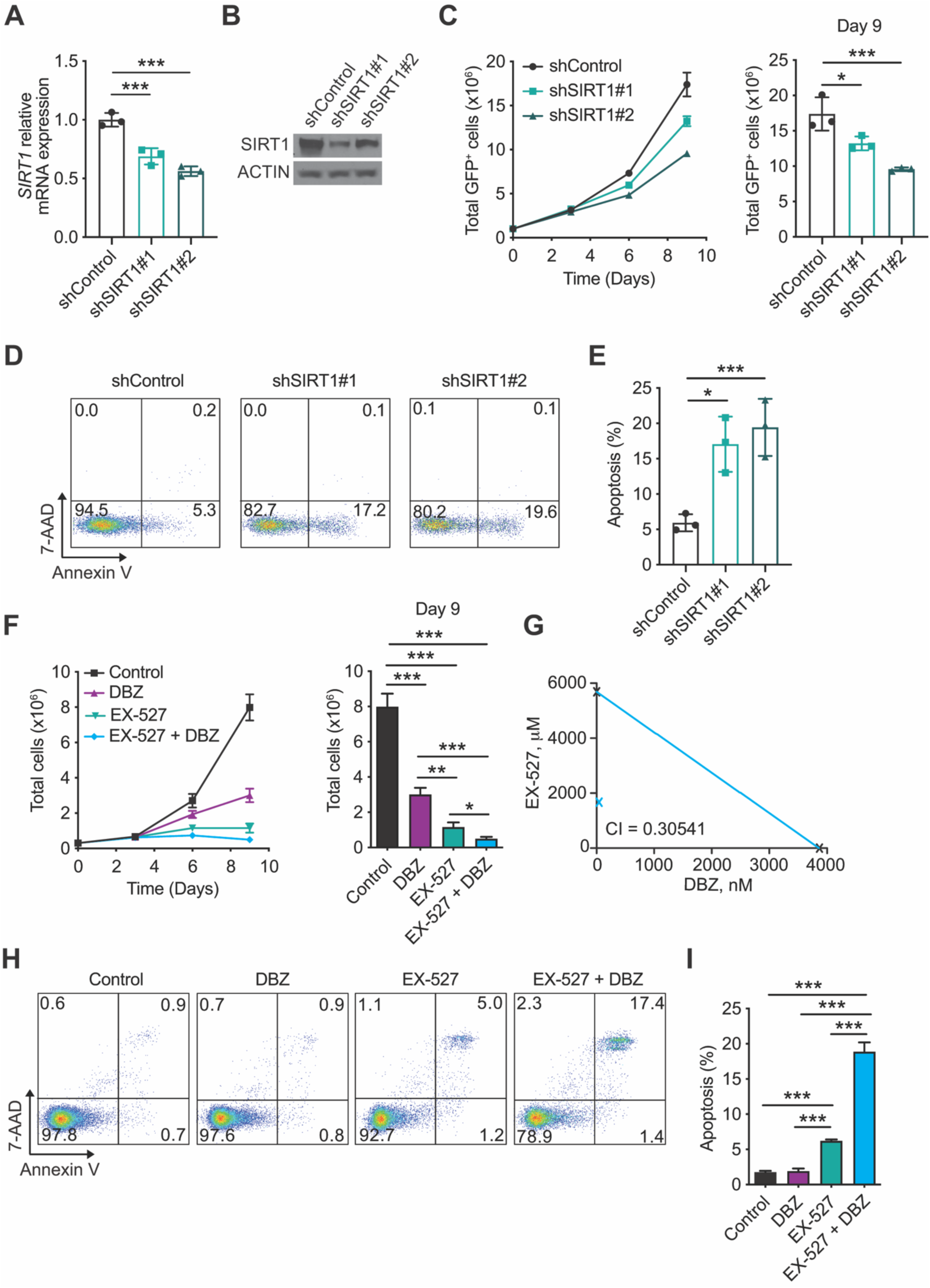
SIRT1 inhibition shows antileukemic and synergistic effects with NOTCH1 inhibition. **A-B,** *SIRT1* mRNA expression levels (**A**) and protein expression levels (**B**) in DND41 cells harboring two independent doxycycline-inducible shRNAs targeting *SIRT1* with concomitant GFP expression or a non-targeting shRNA control, 3 days after doxycycline induction. **C,** Proliferation curve (left) and cell quantification at day 9 (right) of DND41 cells upon Doxycycline-induced expression of a control shRNA or shRNAs targeting *SIRT1*. **D-E**, Representative flow cytometry plots from triplicate samples of annexin V (apoptotic cells) and 7-AAD (dead cells) staining (**D**) and quantification of apoptosis (**E**) of DND41 cells 9 days after Doxycycline-induced expression of a control shRNA or shRNAs targeting *SIRT1*. Numbers in quadrants indicate percentage of cells. **F,** Proliferation curve (left) and cell quantification at day 9 (right) of DND41 cells treated with vehicle (DMSO), EX-527 (90μM), DBZ (250nM), or EX-527 and DBZ in combination. **G**, Isobologram analysis of DBZ and EX-527 treatment after 6 days in DND41 cells. The value for the combination index at ED50 is marked in blue. The ED50 for each drug is marked in black. **H-I**, Representative flow cytometry plots from triplicate samples of annexin V (apoptotic cells) and 7-AAD (dead cells) staining (**H**) and quantification of apoptosis (**I**) of DND41 cells treated with vehicle (DMSO), EX-527(90μM), DBZ (250nM) or EX-527 and DBZ in combination at Day 9. Numbers in quadrants indicate percentage of cells. \**P* < 0.05 and \*\*\**P* < 0.005 in Figs. 1A, 1D and 1F using one-way analysis of variance (ANOVA). \**P* < 0.05 \*\**P* < 0.01 and \*\*\**P* < 0.005 in Figs. 1H and 1K using two-way ANOVA for multiple comparisons.

Moreover, we tested the effects of pharmacologically inhibiting SIRT1 using EX-527, a well-described SIRT1 specific inhibitor (65), alone or in combination with the GSI DBZ. Importantly, EX-527 treatment in DND41 cells impaired cell growth (**Fig. 2F**) and showed strong and synergistic antileukemic activity in combination with DBZ (combination index=0.3 using isobologram analyses) (**Fig. 2G**). Similar to genetic inhibition of *SIRT1*, this effect was primarily driven by increased cytotoxicity, as revealed by increased AnnexinV-positive apoptotic cells (**Fig. 2H-I**). Overall, our results demonstrate that targeting SIRT1 either via genetic or pharmacological means leads to antileukemic and synergistic effects with NOTCH1 inhibition *in vitro*.

### SIRT1 promotes T-ALL development and confers resistance to NOTCH1 inhibition in a deacetylase-dependent manner

Following up on these results, we tested the effects of different *Sirt1* allelic dosages in NOTCH1-induced T-ALL generation. To this end, we generated NOTCH1-induced leukemias in mice as previously described by using retroviral expression of an oncogenic version of NOTCH1 (E-NOTCH1, lacking NOTCH1 extracellular domain) in bone marrow progenitor cells followed by transplantation into irradiated mice (15). In this context, NOTCH1-induced leukemias generated from *Sirt1*-overexpressing bone marrow progenitors (48) showed significantly accelerated kinetics of leukemia development as compared to those generated from *Sirt1* wild-type mice (**Fig. 3A-C**). Conversely, tamoxifen-induced deletion of *Sirt1* only two days after transplantation of *Sirt1* conditional knockout bone marrow progenitor cells infected with an oncogenic form of NOTCH1 resulted in delayed leukemia generation and reduced disease penetrance (**Fig. 3D-F**). Finally, we postulated that SIRT1 might also contribute to mediate resistance to NOTCH1 inhibition *in vivo*. To test this hypothesis, we overexpressed *Sirt1* in already established mouse primary GSI-sensitive NOTCH1-induced leukemias and analyzed its effects on the response to GSI treatment, as previously described (15). In this context, mice were transplanted with a combination of non-transduced (GFP-only) or transduced (GFP/mCherry double positive) T-ALL cells with either the *Sirt1* wild-type deacetylase or a catalytically-dead, H355A-mutant version of *Sirt1* (25) (**Fig. 3G**). Mice transplanted with these T-ALL cells were then treated with the DBZ GSI on a daily basis and leukemia progression was analyzed in peripheral blood (**Fig. 3G**). Notably, while non-transduced GFP-only cells showed little or no progression under treatment, T-ALL cells overexpressing *Sirt1* did progress under GSI treatment *in vivo* (**Fig. 3H-I**). Importantly, this effect was mediated by SIRT1 deacetylase activity since overexpression of the catalytically dead, H355A-mutant version resulted in abrogation of this competitive advantage (**Fig. 3H-I**). Together, our results demonstrate that SIRT1 promotes NOTCH1-induced T-ALL development and mediates resistance to NOTCH1 inhibition *in vivo* in a deacetylase-dependent manner.

**Figure 3.**
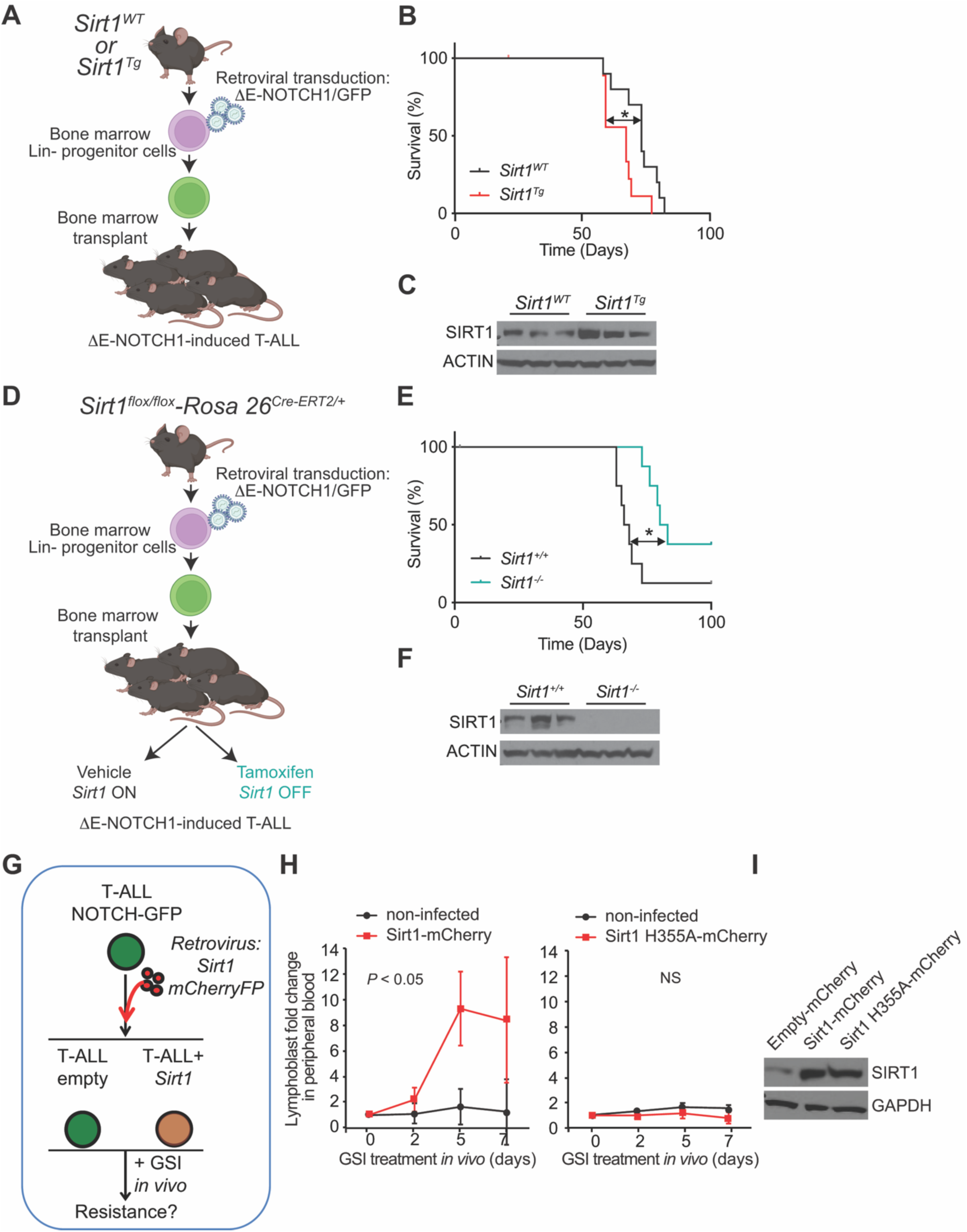
SIRT1 promotes T-ALL development and confers resistance to NOTCH1 inhibition *in vivo*. **A,** Schematic of retroviral-transduction protocol for the generation of NOTCH1-induced T-ALLs from *Sirt1*-overexpressing (48) (*Sirt1*^TG^) or wild-type control littermate (*Sirt1*^WT^) mice. **B,** Kaplan-Meier curves of mice transplanted with E-NOTCH1 infected *Sirt1*^WT^ and *Sirt1*^TG^ hematopoietic progenitors (n=10 per genotype). \**P* < 0.05 value was calculated using the log-rank test. **C,** Western blot analysis of SIRT1 and ACTIN expression in leukemic spleens from terminally ill mice from survival curve in B. **D,** Schematic of retroviral-transduction protocol for the generation of NOTCH1-induced T-ALLs from inducible *Sirt1*-conditional knockout mice. Two days upon transplantation of NOTCH1-infected *Sirt1*^flox/flox^-Rosa26^Cre-ERT2/+^ progenitors, mice were treated with corn oil vehicle (*Sirt1*^+/+^) or tamoxifen (*Sirt1*^-/-^), in order to induce isogenic loss of *Sirt1*. **E,** Kaplan-Meier curves of mice transplanted with NOTCH1-infected *Sirt1*^flox/flox^-Rosa26^Cre-ERT2/+^ progenitors (n=10 and treated with vehicle (*Sirt1*^+/+^) or tamoxifen (*Sirt1*^-/-^) as in D (n=10 per genotype). \**P* < 0.05 value was calculated using the log-rank test. **F,** Western blot analysis of SIRT1 and ACTIN expression in leukemic spleens from terminally ill mice from survival curve in E. **G,** Schematic for transduction of either wild-type *Sirt1* or a deacetylase-dead H355A *Sirt1* mutant concomitantly expressing the mCherry fluorescent protein in NOTCH1-induced GFP+ primary T-ALL cells followed by transplantation into mice, which were subsequently treated daily with DBZ *in vivo*. **H,** Peripheral blood leukemia infiltration in mice harboring NOTCH1-induced T-ALL cells expressing mCherry and *Sirt1* wild-type or H355A-mutant upon continuous daily treatment with DBZ. Changes in leukemia cell counts of noninfected (mCherry-negative) cells are shown as an internal control. Error bars, median ± s.d.; *P* values were calculated using two-tailed Student *t*-test (n=5 mice per group); NS, not significant. **I, C,** Western blot analysis of SIRT1 and GAPDH expression in leukemic spleens from non-DBZ-treated, terminally ill mice from G.

### Secondary loss of SIRT1 in established T-ALL shows antileukemic and highly synergistic effects with NOTCH1 inhibition in vivo

We next investigated the therapeutic effects of targeting *Sirt1* in already established NOTCH1-induced leukemias *in vivo*. To this end, we followed the same retroviral-transduction protocol described before using bone marrow cells from *Sirt1* conditional knockout mice, but we first let leukemias fully develop before analyzing the effects of secondary SIRT1 loss (**Fig. 4A**). Here, we tested the effects of SIRT1 loss not only in leukemias driven by a E-NOTCH1 construct, but also in leukemias driven by a weaker oncogenic version of NOTCH1 that harbors the prototypical mutations targeting the NOTCH1 heterodimerization domain (HD) and the PEST domain (HD P-NOTCH1), which are recurrently observed in patient samples (9) (**Fig. 4A**). In this context, tamoxifen-induced isogenic deletion of *Sirt1* in already established primary *Sirt1* conditional knockout leukemias led to significant and highly synergistic antileukemic effects with GSI treatment *in vivo*, regardless of the oncogenic version of NOTCH1 used (**Fig. 4B**). Interestingly, leukemias that eventually relapsed after tamoxifen injection did show SIRT1 loss (**Fig. 4C**), suggesting that T-ALL cells can still adapt to survive *Sirt1* deletion. To further dissect the acute effects of SIRT1 loss in T-ALL *in vivo*, we transplanted the same E-NOTCH1 *Sirt1* conditional knockout leukemias in mice and let them become overtly leukemic before inducing SIRT1 loss. In this setting, analyses of leukemic spleens only 48h after tamoxifen injection revealed drastically reduced SIRT1 levels (**Fig. 4D-E**), together with reduced tumor burden (**Fig. 4F-G**) at the expense of increased apoptosis (**Fig. 4H-I**). Similar effects were observed upon acute loss of SIRT1 in mice harboring HD P-NOTCH1 *Sirt1* conditional knockout leukemias (**Supplementary Fig. S2**). Overall, our results demonstrate the antileukemic and highly synergistic effects with NOTCH1 inhibition of genetically targeting *Sirt1* in T-ALL *in vivo*.

**Figure 4.**
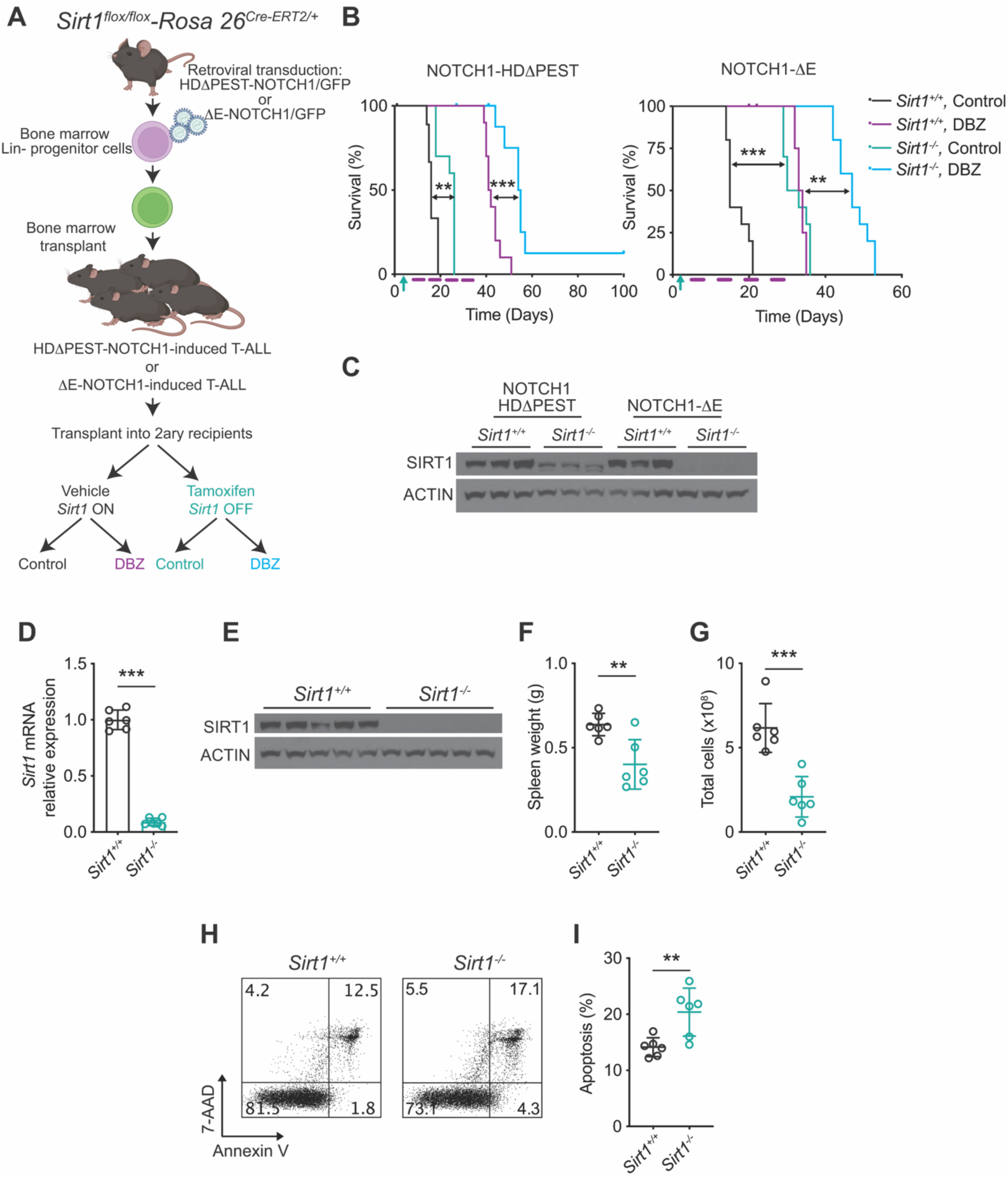
Secondary loss of SIRT1 in established leukemias leads to antileukemic and synergistic effects with NOTCH1 inhibition *in vivo*. **A,** Schematic of retroviral-transduction protocol for the generation of NOTCH1-induced T-ALLs from inducible *Sirt1*-conditional knockout mice, followed by transplant into secondary recipients treated with vehicle (*Sirt1*^+/+^) or tamoxifen (*Sirt1*^-/-^) and vehicle or DBZ. **B,** Kaplan-Meier survival curves of mice harboring *Sirt1*-positive and *Sirt1*-deleted isogenic leukemias treated with 4 cycles of vehicle or DBZ (5 mg/kg) on a 4-days-ON (red blocks) and 3-days-OFF schedule (log-rank test; \*\**P* < 0.01; \*\*\**P* < 0.005; n = 10 per group). **C,** Western blot analysis of SIRT1 and ACTIN expression in leukemic spleens from terminally ill mice from survival curve in B. **D-E**, Quantitative RT-PCR analysis of *Sirt1* mRNA expression (D) and western blot analysis of SIRT1 protein levels (E) in tumor cells isolated from E-NOTCH1-induced *Sirt1* conditional knockout leukemia–bearing mice 48 h after being treated with vehicle only (*Sirt1*^+/+^) or tamoxifen (*Sirt1*^-/-^) *in vivo*. **F-G,** Tumor burden in E-NOTCH1-induced *Sirt1* conditional knockout leukemia–bearing mice 48 h after being treated with vehicle only (*Sirt1*^+/+^) or tamoxifen (*Sirt1*^-/-^) *in vivo* as revealed by total spleen weight (F) and total spleen cell numbers (G). **H-I**, Representative flow cytometry plots from of annexin V (apoptotic cells) and 7-AAD (dead cells) staining (**H**) and quantification of apoptosis (**I**) in leukemic spleens from E-NOTCH1-induced *Sirt1* conditional knockout leukemia– bearing mice 48 h after being treated with vehicle only (*Sirt1*^+/+^) or tamoxifen (*Sirt1*^-/-^) *in vivo*. (n = 5 per treatment; \*\**P* < 0.01 and \*\*\**P* < 0.005 in Figs. 1D-I using two-tailed Student *t*-test).

### Acute loss of SIRT1 leads to AMPK activation and global metabolic effects

SIRT1 is a well-known metabolic regulator and SIRT1 loss has been shown to impact metabolism in other hematological malignancies (40); thus, we next investigated the metabolic effects of its loss. In addition to increased acetylation levels of p53 and p65 NFκB, two well-described SIRT1 targets (**Supplementary Fig. S3A-B**), we observed activation of AMPK and downregulation of mTOR signaling as revealed by decreased p4E-BP1 levels, together with reduced levels of the stress response factor ATF4 (**Fig. 5A** and **Supplementary Fig. S3C**), suggesting SIRT1 loss results in a metabolic crisis in leukemia cells. We next performed global untargeted metabolomic analyses (LC-MS) upon acute loss of SIRT1 in T-ALL *in vivo*. These experiments revealed that SIRT1 loss leads to global metabolic changes (**Fig. 5B**), including accumulation of glycolytic intermediates (**Fig. 5C**), together with increased levels of glutamine, and glutamine-derived metabolites such as glutamate, aspartate and asparagine (**Fig. 5D**). In addition, seahorse analyses in *Sirt1* conditional knockout leukemia-derived cell lines revealed a global reduction in oxygen consumption rate (OCR) upon SIRT1 loss (**Fig. 5E** and **Supplementary Fig. S3D**). Using these murine tumor-derived cell lines, we also observed synergistic effects with NOTCH1 inhibition *in vitro* driven by increased cytotoxicity (**Supplementary Fig. S3E-H**), consistent with the effects previously observed in human cells (**Fig. 2**).

**Figure 5.**
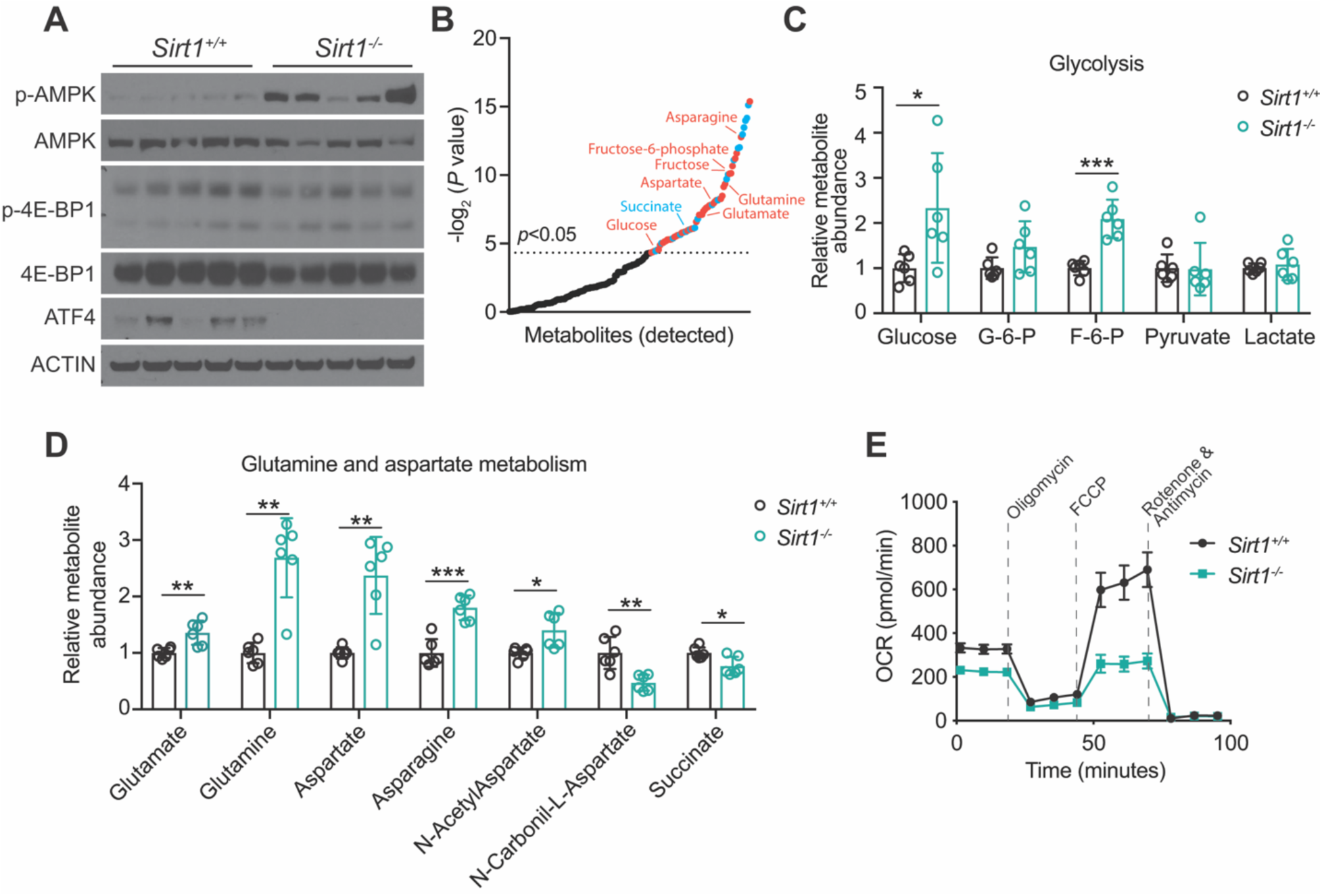
Metabolic consequences of secondary loss of SIRT1 in established leukemias *in vivo*. **A,** Western blot analysis of AMPK protein levels and activation, 4E-BP1 levels and activation, and ATF4 levels in tumor cells isolated from E-NOTCH1-induced *Sirt1* conditional knockout leukemia–bearing mice 48 h after being treated with vehicle only (*Sirt1*^+/+^) or tamoxifen (*Sirt1*^-/-^) *in vivo*. **B,** Significantly altered metabolites (upregulated in red, downregulated in blue) upon tamoxifen-induced isogenic loss of *Sirt1* in leukemic spleens from mice treated as in A, ranked by *P* value (–log10 transformed). **C-D,** Relative abundance of indicated glycolytic intermediates (C) or glutamine/aspartate-related metabolites (D) upon tamoxifen-induced isogenic loss of *Sirt1* in leukemic spleens from mice treated as in A. (n = 5 per treatment; \**P* < 0.05, \*\**P* < 0.01 and \*\*\**P* < 0.005 in Figs. 5A-D using two-tailed Student *t*-test). **E,** Oxygen consumption rate (OCR) in response to the indicated mitochondrial inhibitors in a E-NOTCH1-induced *Sirt1* conditional knockout leukemia-derived cell line under basal conditions or 2-days after 4-Hydroxytamoxifen-induced isogenic loss of *Sirt1*, measured in real time using a Seahorse XF24 instrument. Data are presented as +/- SD of n = 5 wells.

We next asked whether the strong AMPK activation observed might be mediating the therapeutic effects of SIRT1 loss. To address this question, we generated double *Sirt1*/*Ampk* conditional knockout leukemias following the same approach previously described (**Supplementary Fig. S4A**). However, tamoxifen-induced secondary loss of SIRT1 in combination with AMPK still resulted in significant antileukemic effects with extended survival (**Supplementary Fig. S4B-C**), suggesting that AMPK activation is dispensable for the therapeutic effects of SIRT1 loss. Similarly, given the global metabolic changes affecting glycolysis and glutaminolysis, we hypothesized that metabolic rescues using membrane soluble compounds that can bypass glycolysis or glutaminolysis blocks, such as methyl-pyruvate (MP) or dimethyl-2-oxoglutarate (DMKG), might rescue the effects of SIRT1 loss, similar to what we previously described upon NOTCH1 inhibition (15). Yet, addition of either MP or DMKG did not prevent the antileukemic effects and impaired proliferation upon SIRT1 loss *in vitro* (**Supplementary Fig. S4D-E**). Overall, these data shows that SIRT1 loss leads to AMPK activation and global metabolic changes although these might be dispensable for its therapeutic effects.

### Acute loss of SIRT1 results in KAT7 hyperacetylation, a transcriptional signature driven by KAT7 loss and globally decreased H4K12ac levels

Given the deacetylase-dependent effects of SIRT1 on the resistance to NOTCH1 inhibition (**Fig. 3H-I**), we next postulated that unbiased acetyl-proteomic experiments might reveal relevant targets for the mechanistic effects of SIRT1 loss in T-ALL. To this end, we performed Mass Spectrometry analyses upon Acetyl-Lysine enrichment in *Sirt1* conditional knockout T-ALLs treated acutely with tamoxifen or vehicle control *in vivo* (**Fig. 6A**). These experiments revealed multiple hyperacetylated peptides upon SIRT1 loss (**Supplementary Table S1**). Interestingly, integrative results using our two independent *Sirt1* conditional knockout leukemias revealed only nine targets that were consistently hyperacetylated (**Fig. 6A**). These notably included hyperacetylation of histone acetyltransferase 7 (KAT7, also known as HBO1) on lysine 277, without affecting total KAT7 levels (**Fig. 6A** and **Supplementary Table S1**); we also detected hyperacetylation of Bromodomain-containing protein 1 (BRD1) on Lys 418. Since both KAT7 and BRD1 are known to form part of a histone acetyltransferase complex which targets H4K12 and H3K14 for acetylation (66), our results suggested that SIRT1 loss might be affecting the activity of this complex. Therefore, we performed gene expression profiling upon acute loss of SIRT1 in T-ALL *in vivo*. While these analyses revealed global transcriptional changes upon SIRT1 loss (**Fig. 6B** and **Supplementary Table S2**), gene set enrichment analyses (GSEA) against the C2 database showed no evident signature related to glycolysis, glutaminolysis, oxidative phosphorylation and/or PGC1α as significantly enriched upon *Sirt1* deletion (**Supplementary Table S3**). Recent literature has shown KAT7 loss leads to antileukemic effects in MLL-rearranged leukemias (56). Thus, we tested our transcriptional signature after loss of SIRT1 against those observed upon KAT7 loss (56) and uncovered a significant correlation between both (**Fig. 6C**), suggesting that SIRT1 loss leads to hyperacetylation of KAT7, which might be less active. Consistent with this hypothesis, we detected a global reduction in H4K12ac upon acute loss of SIRT1 in mouse T-ALL *in vivo* (**Fig. 6D**). Moreover, doxycycline inducible knockdown of *SIRT1* in human DND41 T-ALL cells led to similarly reduced levels of H4K12ac (**Fig. 6E**). Further ChIPseq epigenetic profiling of H14K12ac upon deletion of *Sirt1* confirmed a global reduction in H4K12ac peaks *in vivo* (**Fig. 6F** and **Supplementary Table S4**). In addition, integration of these data with our gene expression profiling data revealed that regions showing reduced H4K12ac levels were significantly enriched in the promoter of downregulated genes upon SIRT1 loss (**Fig. 6G** and **Supplementary Fig. S5**). Together, our data demonstrates that SIRT1 loss leads to hyperacetylation of KAT7, which correlates with the transcriptional effects of KAT7 loss and results in globally decreased levels of the KAT7 mark H4K12ac.

**Figure 6.**
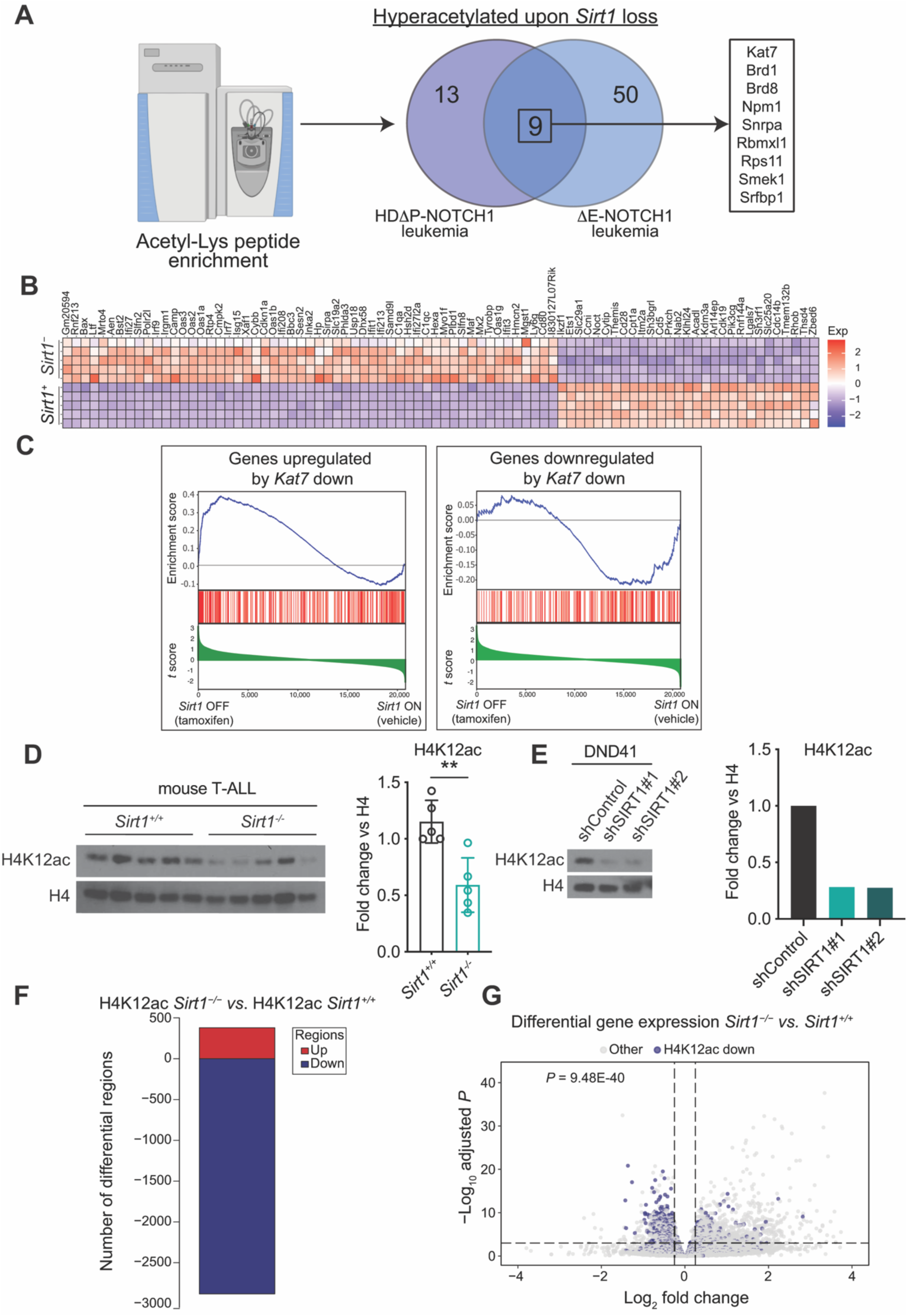
Secondary loss of SIRT1 leads to hyperacetylation of KAT7 and a transcriptional signature driven by KAT7 inhibition. **A,** Schematic representation of acetyl-proteomic experiments in tumor cells isolated from E-NOTCH1 or HD P-NOTCH1- induced *Sirt1* conditional knockout leukemia–bearing mice 48 h after being treated with vehicle only (*Sirt1*^+/+^) or tamoxifen (*Sirt1*^-/-^) *in vivo*. Venn diagram shows significantly hyperacetylated targets consistently found upon SIRT1 loss in both leukemias. **B,** Heatmap representation of the top differentially expressed genes between control (*Sirt1*^+/+^) and tamoxifen-treated (*Sirt1*^-/-^) *Sirt1* conditional knockout E-NOTCH1–induced leukemias. Cutoffs used: Wald statistic < −8 or > 8; P-adjusted value < 0.005; sorted based on mean expression levels. Scale bar shows color-coded differential expression, with red indicating higher levels of expression and blue indicating lower levels of expression. **C,** GSEA of genes regulated by KAT7 in vehicle only–treated (*Sirt1*^+/+^) compared with tamoxifen–treated (*Sirt1*^-/-^) E-NOTCH1-induced *Sirt1* conditional knockout leukemia cells *in vivo*. **D,** Western blot analysis (left) and quantification (right) of H4 total protein levels and H4K12ac levels in tumor cells isolated from E-NOTCH1- induced *Sirt1* conditional knockout leukemia–bearing mice 2 days after being treated with vehicle only (*Sirt1*^+/+^) or tamoxifen (*Sirt1*^-/-^) *in vivo*. \*\**P* < 0.01 using two-tailed Student *t*-test. **E,** Western blot analysis (left) and quantification (right) of H4 total protein levels and H4K12ac levels in DND41 cells harboring two independent doxycycline-inducible shRNAs targeting *SIRT1* with concomitant GFP expression or a non-targeting shRNA control, 3 days after doxycycline induction. **F,** Bar graph showing the number of significantly downregulated (blue) or upregulated (red) H4K12ac-containing genomic regions from H4K12ac ChIP-seq analyses in tumor cells isolated from E-NOTCH1-induced *Sirt1* conditional knockout leukemia–bearing mice 48 h after being treated with vehicle only (*Sirt1*^+/+^) or tamoxifen (*Sirt1*^-/-^) *in vivo*. **G,** Volcano plot showing H4K12ac downregulated region genes (shrunken LFC < -0.3) in blue (n=2319) and the rest of the regions in light grey. Horizontal dashed line corresponds to the adjusted *P* value threshold of 0.001. Vertical dashed lines correspond to log_2_FC changes of +0.25 and -0.25. 303/2319 were significantly downregulated at the gene expression level (hypergeometric test *P* value = 9.48E-40).

### KAT7 partially mediates the antileukemic effects of SIRT1 loss

Our previous data suggested that KAT7 inactivation might drive, at least in part, the antileukemic effects of SIRT1 loss. Thus, we hypothesized that *Sirt1*-deleted leukemia cells might be more sensitive to KAT7 inhibition than wild-type counterparts. Indeed, experiments using the KAT7-specific inhibitor WM-3835 (67) in our *Sirt1* conditional knockout T-ALL cell line revealed significantly stronger antileukemic effects in *Sirt1*-deleted leukemia cells at different concentrations, resulting in a lower IC50 than in isogenic *Sirt1*-positive leukemia cells (**Fig. 7A**). Finally, to formally test whether KAT7 mediates part of the antileukemic effects of SIRT1 loss, we performed rescue experiments overexpressing either wild-type KAT7, an acetyl-mimic (K277Q) or an acetyl-dead (K277R) version of KAT7 in our *Sirt1* conditional knockout T-ALL cell line (**Fig. 7B**). Notably, while wild-type or K277Q KAT7 did not affect the impaired proliferation upon SIRT1 loss (**Fig. 7C**), overexpression of the acetyl-dead K277R version of KAT7 was able to partially rescue the antileukemic effects of SIRT1 loss, resulting in increased cell proliferation (**Fig. 7C**) and reduced apoptosis (**Fig. 7D**). Overall, these results demonstrate that the antileukemic effects of SIRT1 loss are partly mediated by KAT7.

**Figure 7.**
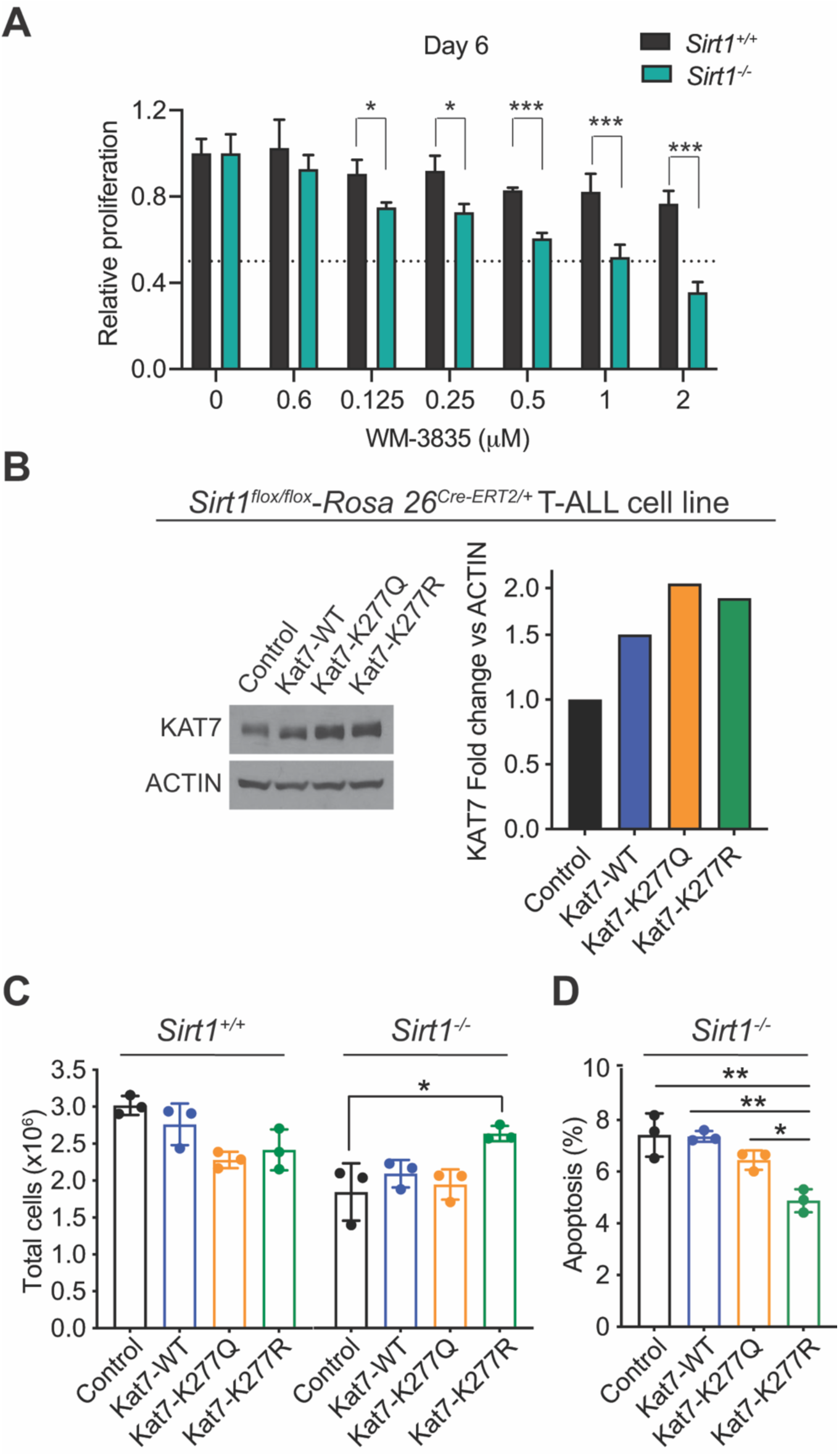
KAT7 partially mediates the antileukemic effects of loss of SIRT1. **A,** Relative cell proliferation of a E-NOTCH1-induced *Sirt1* conditional knockout leukemia-derived cell line upon treatment with ethanol (*Sirt1*^+/+^) or 4-Hydroxytamoxifen (*Sirt1*^-/-^) and different concentrations of the KAT7 inhibitor WM-3835 *in vitro*. \**P* < 0.05 and \*\*\**P* < 0.005 using two-tailed Student *t*-test. **B,** Western blot analysis (left) and quantification (right) of a E-NOTCH1-induced *Sirt1* conditional knockout leukemia-derived cell line infected with an empty vector (control) or with constructs overexpressing wild-type KAT7, K277Q mutant KAT7 or K277R mutant KAT7. **C,** Quantification of total cell numbers 6 days after 4-hydroxytamoxifen-induced loss of SIRT1 in cells overexpressing an empty vector or different KAT7 versions from B. **D,** Quantification of apoptosis (Annexin V-positive cells) 6 days after 4-hydroxytamoxifen-induced loss of SIRT1 in cells overexpressing an empty vector or different KAT7 versions from B. \**P* < 0.05 and \*\**P* < 0.01 in Figs. 7C-D using 2-way analysis of variance (ANOVA) for multiple comparisons.

## Discussion

Here, we demonstrate that *SIRT1* is a direct target of NOTCH1 in T-ALL through a newly identified enhancer. Accordingly, SIRT1 is broadly overexpressed in T-ALL. Genetic experiments *in vivo* demonstrated an oncogenic role for SIRT1 in both leukemia generation and progression such that targeting SIRT1 pharmacologically *in vitro*, or genetically *in vivo*, leads to highly antileukemic and synergistic effects with GSIs. Mechanistically, SIRT1 loss leads to global transcriptional and metabolic changes together with hyperacetylation and inactivation of the KAT7 histone acetyltransferase, which is critical for the therapeutic effects of SIRT1 loss. Consistently, a non-deacetylatable mutant form of KAT7 partially rescues the effects of SIRT1 loss on leukemia proliferation. Overall, we uncovered a prominent role for in T-ALL generation and progression, as well as a novel therapeutic target in T-ALL. In this context, it is relevant to note the multiple efforts and compounds described to date to either pharmacologically activate or inhibit SIRT1 (68). Still, the field of SIRT1 pharmacological modulation has been mired in controversy, with many studies disputing the specificity of these compounds, most prominently resveratrol and other purported SIRT1 activators (69). While described SIRT1 inhibitors are reported to be more specific, none of the inhibitors described to date have been clinically approved, probably owing to their poor bioavailability *in vivo* (70). In this setting, our results demonstrating that SIRT1 inhibition could have relevant roles in the treatment of T-ALL *in vivo* will hopefully spur renewed efforts to discover improved SIRT1 inhibitor drugs for clinical development.

Our finding of a novel NOTCH1-bound *SIRT1* enhancer (N-Se) adds to the growing literature of relevant enhancers in T-ALL, including the N-Me *MYC* enhancer (44), the PE *PTEN* enhancer (47) or the *de novo* acquired enhancers for TAL1 (71) and BCL11B (72), among others (73). Our results unequivocally demonstrate that N-Se regulates *SIRT1* levels. Still, the binding of NOTCH1 to N-Se is relatively weak compared to N-Me or other enhancer regions, and NOTCH1 inhibition results in a modest reduction of SIRT1 protein levels. Thus, it is likely that other transcription factors might be equally or even more relevant in regulating the effects of N-Se. Further studies to mechanistically dissect the role and mechanisms controlling N-Se activity will answer these questions.

Mechanistically, acute loss of SIRT1 resulted in broad metabolic, transcriptional, and epigenetic changes. From a metabolic point of view, we observed accumulation of glycolytic and glutamine-derived metabolites upon SIRT1 loss, together with strong activation of AMPK, suggesting a metabolic crisis. This is consistent with previous literature describing SIRT1 activates AMPK in a feedforward positive loop (29, 30). As expected, AMPK activation resulted in downregulation of mTOR signaling together with a reduction in ATF4 levels, consistent with previous findings demonstrating mTOR directly activates ATF4 (74). We also observed a concomitant decrease in OCR, similar to results previously described in CML models with loss of SIRT1 (40). Thus, we decided to formally test the relevance of these metabolic effects on the therapeutic effects of SIRT1 loss. However, experiments *in vivo* using double *Sirt1*/*Ampk* conditional knockout leukemias still showed significantly extended survival even after concomitant loss of SIRT1 and AMPK. We previously demonstrated that isogenic loss of AMPK alone does not impact leukemia progression by itself (75). Thus, since the extension in survival observed upon SIRT1/AMPK loss was similar to the one observed upon SIRT1 loss alone, our results suggest that AMPK activation is dispensable for the survival extension upon loss of SIRT1. In addition, the decrease in OCR upon SIRT1 loss in CML models has been mechanistically linked to inhibition of PGC1α (40); yet, our gene expression profiling data did not show any significant correlation with an inactivation of PGC1α transcriptional program, suggesting that other mechanisms might mediate these effects in T-ALL. Finally, experiments using membrane soluble metabolites that bypass blocks in glycolysis and glutaminolysis failed to rescue the impaired proliferation observed upon SIRT1 loss. Together, these results suggest that, although the metabolic effects of SIRT1 loss may play an important role, they are likely not the main mechanistic mediators of its phenotype in T-ALL and, thus, they might be more a consequence rather than a cause of its antileukemic effects.

To gain further insights into the mechanistic effects of SIRT1 loss, we performed global unbiased acetyl-proteomic analyses, together with gene expression and epigenetic profiling in our *Sirt1* conditional leukemias. While we did not observe differences in the acetylation levels of previously described SIRT1 targets such as NOTCH1 or MYC, our analyses revealed KAT7 and BRD1 as consistently hyperacetylated upon SIRT1 loss. This finding caught our attention since KAT7 and BRD1 are known to form part of the same histone acetyltransferase complex targeting H4K12 and H3K14 (66). Moreover, KAT7 has been recently shown to play important roles in AML (56,67,76), which suggests that its hyperacetylation might be involved in the mechanistic therapeutic effects of SIRT1 loss. Indeed, gene expression profiling upon SIRT1 loss in T-ALL *in vivo* revealed a transcriptional signature similar to the one observed upon KAT7 loss. Consistently, we observed globally reduced levels of H4K12ac upon SIRT1 loss in mouse or human T-ALL cells. Finally, overexpression rescue experiments in our *Sirt1* conditional knockout leukemia-derived cell line using either wild-type KAT7 or acetyl-mimic (K277Q) or acetyl-dead (K277R) KAT7 mutant versions showed that overexpression of the non-acetylatable K277R KAT7 partially rescued the antiproliferative and cytotoxic effects of SIRT1 loss. Overall, our experiments demonstrate that KAT7 mediates, at least in part, the antileukemic effects of SIRT1 loss.

Previous studies have shown that KAT7 is important for the recruitment and processivity of RNApolII (67). Thus, it is tempting to speculate that this effect might be relevant for the broad transcriptional changes observed upon SIRT1 loss. Similarly, KAT7 loss has been shown to impair leukemic stem cell activity in AML (67), as well as to lead to exhaustion of hematopoietic stem cells (76). Interestingly, SIRT1 itself has been shown to play a role in hematopoietic stem cells and leukemic stem cell maintenance in AML (39,40,77), thus, it is also fair to speculate that these phenotypes might be mediated by KAT7. Still, further studies are warranted to fully understand the specific effects of KAT7 in T-ALL, as well as the relevance of its acetylation.

Our results demonstrate that KAT7 partly mediates the effects of SIRT1 loss in T-ALL, still, the proliferative rescue observed is not total. This might be partly due to the high levels of endogenous KAT7 present in leukemia cells, which might obscure the effects of additional wild-type/mutant KAT7 overexpression. In addition, it may be possible that concomitant overexpression of acetyl-dead mutants for both KAT7 and BRD1 is needed to completely rescue the effects of SIRT1 loss, since they are both involved in the same histone acetyltransferase complex. It is also possible that some of the other seven consistently hyperacetylated proteins upon loss of SIRT1 might play relevant roles and partly influence the antileukemic effects of *Sirt1* deletion. Among these, the hyperacetylation in NPM1 was particularly interesting, given the well-described roles of *NPM1* mutations in AML (78). Still, nothing is known regarding a putative role for NPM1 in T-ALL, thus, further studies are needed to clarify the potential relevance of these findings.

Finally, SIRT1 has already been described to play pleiotropic roles in the literature, where it has been shown to affect the acetylation of multiple targets with relevant roles in leukemia. While we did not detect increased acetylation of NOTCH1 or MYC in our mass spectrometry data, we detected an expected increase in the acetylation of p53 upon SIRT1 loss, which is known to activate p53 (35, 36). Thus, it is possible that part of the antileukemic effects observed upon SIRT1 loss might be mediated by p53. Still, we observed antileukemic effects for genetic or pharmacological inhibition of SIRT1 in human DND41 T-ALL cells, which are known to be p53 mutant (79), suggesting that p53 might not be the central factor mediating the observed effects. Overall, our data revealed wide-ranging effects upon SIRT1 loss in leukemia and demonstrated that part of its antileukemic effects are mediated by the hyperacetylation and inhibition of KAT7.

Finally, our data uncovered a novel feedback mechanism balancing the activity of histone deacetylases and acetyl-transferases. Our results demonstrate that reduced activity of the SIRT1 histone deacetylase is paralleled by a concomitant increase in the acetylation of KAT7, resulting in inhibition of its acetyltransferase activity. Thus, a decrease in the histone deacetylase activity of SIRT1 would be paralleled by a decrease in KAT7 histone acetyltransferase activity. Conversely, an increase of SIRT1 histone deacetylase activity should be paralleled by reduced KAT7 acetylation together with an increase its histone acetyltransferase activity. In this way, the interplay between SIRT1 and KAT7 constitutes a novel rheostat mechanism regulating the total epigenetic output in terms of histone acetylation levels, which is reminiscent of other well-described relevant feedback mechanisms, such as the tightly regulated interplay between protein kinases and phosphatases in mitosis and beyond (80–82). Thus, while our data unveils a SIRT1-KAT7 link with clear relevance in T-cell leukemia, our results point to the existence of deacetylase-histone acetyltransferase balancing mechanisms which might be broadly relevant across different biological processes and cancer types.

Overall, our results reveal an oncogenic role for SIRT1 in T-ALL generation and progression downstream of NOTCH1, identify SIRT1 as a novel therapeutic target for the treatment of T-ALL and uncover a novel SIRT1-KAT7 link that might represent a more broadly relevant mechanism in cancer.

## Acknowledgements

We are grateful to Adolfo A. Ferrando (Columbia University Medical Center) for sharing with us normal human thymocyte samples, and we thank both Adolfo A. Ferrando and Antonio Maraver (IRCM, Montpellier) for their constant constructive criticism and support. We also thank everyone involved with JuanLord for their support. Figures 3A, 3D, 4A and 6A were created using BioRender.com.

## Author contributions

O.L. performed most molecular biology, cellular and animal experiments. A.S. performed all computational analyses. V.dS., L.T., P.R.N., J.K., Z.Z and P.P.R. assisted with molecular experiments. M.A. and S.L. assisted with mouse experiments. C.Z. and H.Z. performed and analyzed acetyl-protemics data. E.C. assisted with metabolomic experiments. X.S. supervised metabolomic analyses. H.K. supervised all computational analyses. D.H. designed the study, supervised the research and wrote the manuscript with O.L., with input from all authors.

