## Supplementary Figures for "A therapeutically targetable NOTCH1-SIRT1-KAT7 axis in T-cell Leukemia"

**Supplementary Figures and Figure Legends**

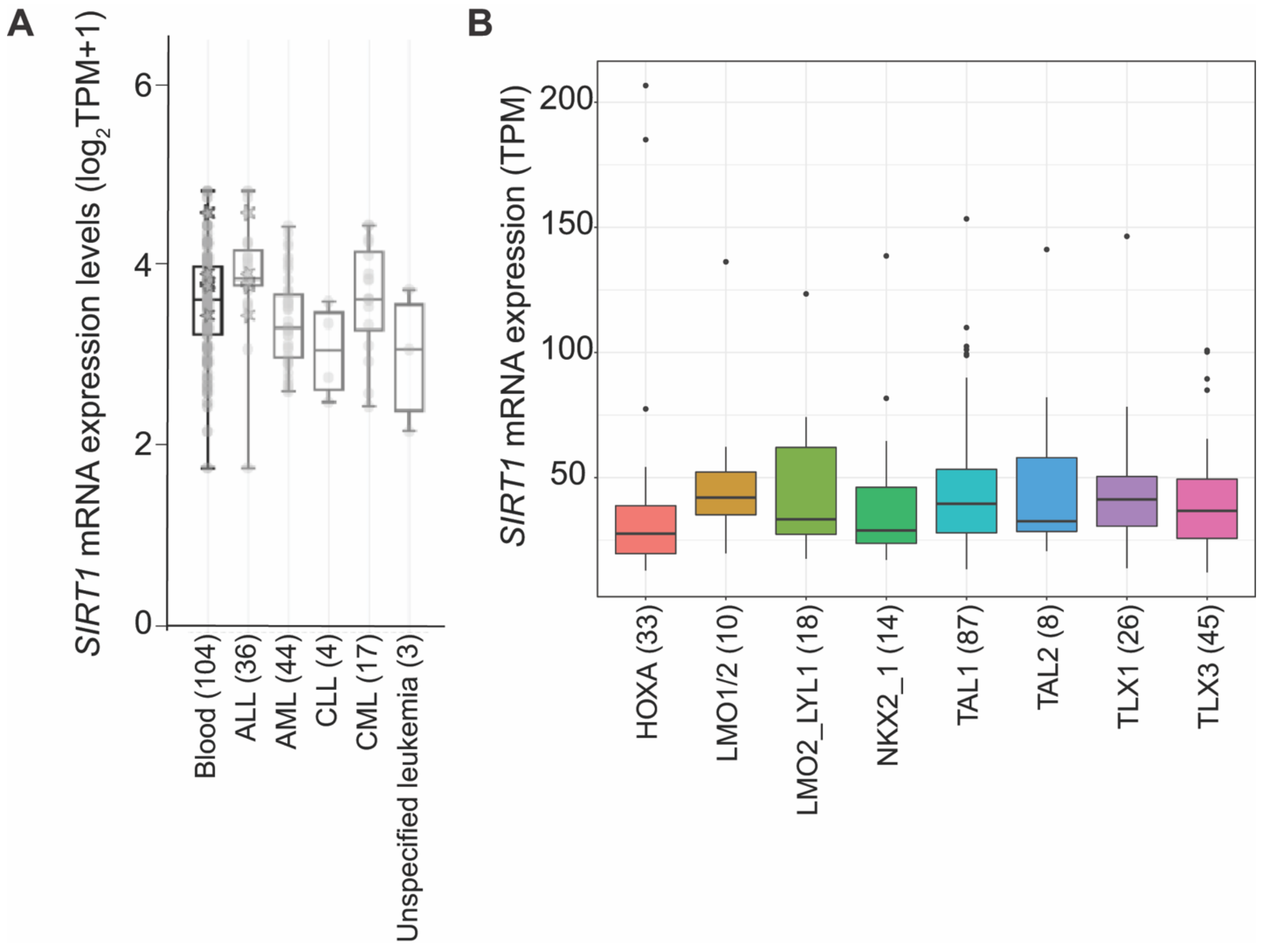


**Supplementary Figure S1**. *SIRT1* expression in human T-ALL. **A,** Box-plots showing *SIRT1* expression in T-ALL cell lines as compared to cell lines from other hematological malignancies available from the CCLE database (62). **B,** *SIRT1* expression levels in human primary T-ALL cases from different subclinical groups (63).

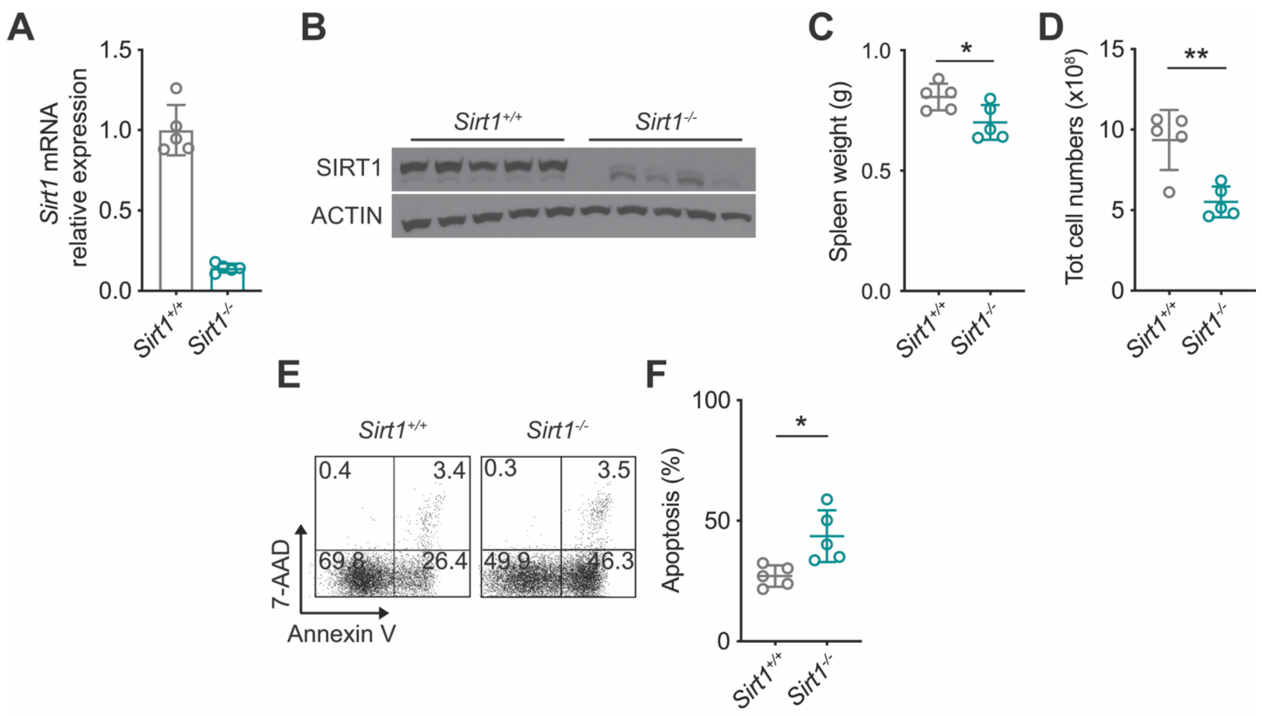


**Figure 2.** Secondary loss of SIRT1 in HDΔP-NOTCH1-induced T-ALL shows antileukemic effects. **A-B**, Quantitative RT-PCR analysis of *Sirt1* mRNA expression (A) and western blot analysis of SIRT1 protein levels (B) in tumor cells isolated from HDΔP-NOTCH1-induced *Sirt1* conditional knockout leukemia–bearing mice 48 h after being treated with vehicle only (*Sirt1*$^{+/+}$) or tamoxifen (*Sirt1*$^{-/-}$) *in vivo*. **C-D,** Tumor burden in HDΔP-NOTCH1-induced *Sirt1* conditional knockout leukemia–bearing mice 48 h after being treated with vehicle only (*Sirt1*$^{+/+}$) or tamoxifen (*Sirt1*$^{-/-}$) *in vivo* as revealed by total spleen weight (C) and total spleen cell numbers (D). **E-F**, Representative flow cytometry plots from of annexin V (apoptotic cells) and 7-AAD (dead cells) staining (**E**) and quantification of apoptosis (**F**) in leukemic spleens from HDΔP-NOTCH1-induced *Sirt1* conditional knockout leukemia–bearing mice 48 h after being treated with vehicle only (*Sirt1*$^{+/+}$) or tamoxifen (*Sirt1*$^{-/-}$) *in vivo*. (n = 5 per treatment; *$P < 0.05$ and **$P < 0.01$ using two-tailed Student *t*-test).

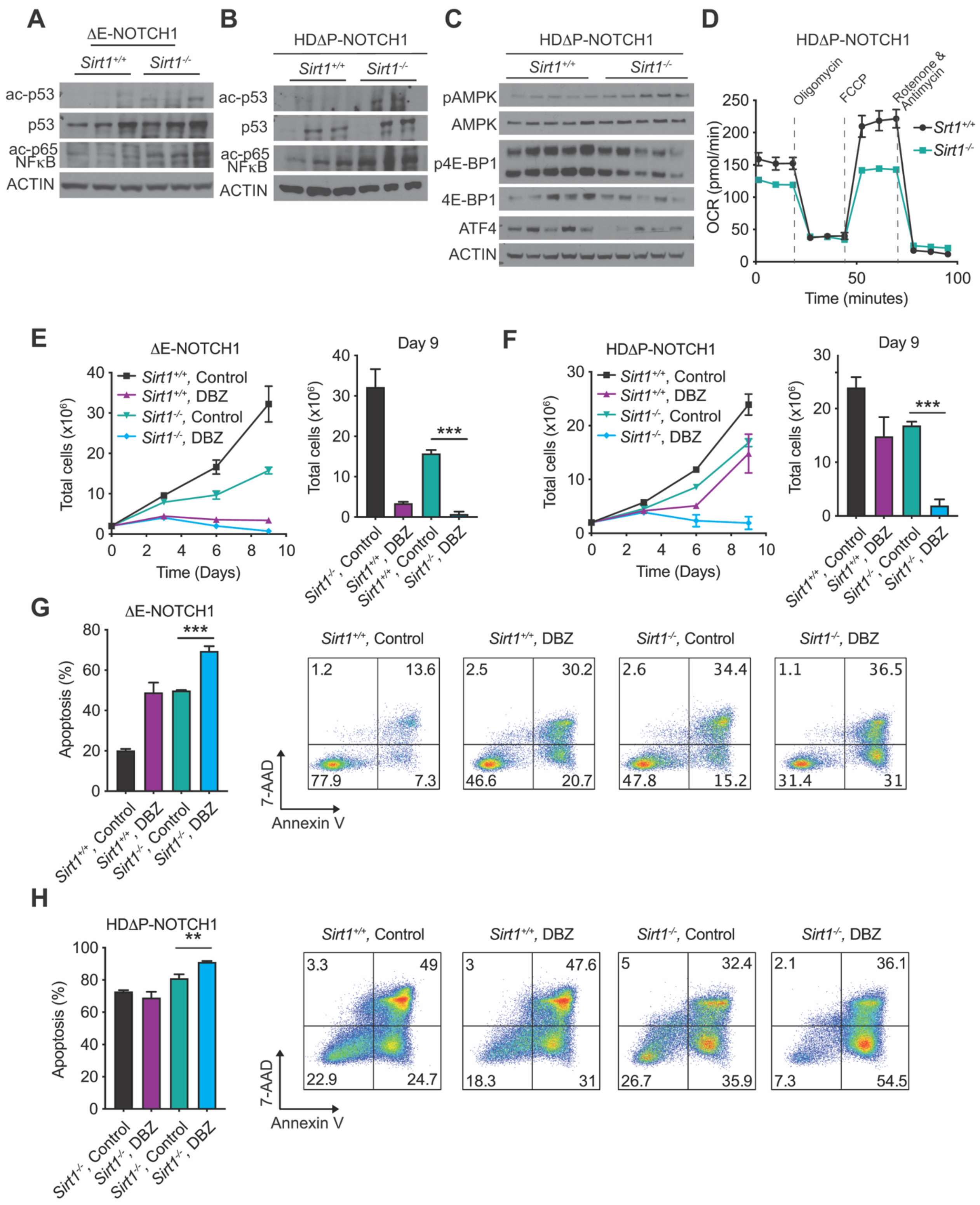


**Figure 3.** Secondary loss of SIRT1 in mouse leukemias *in vivo* leads to hyperacetylation of p53, activation of AMPK, and synergizes with NOTCH1 inhibition *in vitro*. **A-B,** Western

blot analysis of p53 and p65 in tumor cells isolated from ΔE-NOTCH1-induced (A) or HDΔP-NOTCH1-induced (B) *Sirt1* conditional knockout leukemia–bearing mice 48 h after being treated with vehicle only or tamoxifen *in vivo*. **C,** Western blot analysis of AMPK, 4EBP1 and ATF4 in tumor cells isolated from HDΔP-NOTCH1-induced *Sirt1* conditional knockout leukemia–bearing mice 48 h after being treated with vehicle only or tamoxifen *in vivo*. **D,** Oxygen consumption rate (OCR) in response to the indicated mitochondrial inhibitors in a HDΔP-NOTCH1-induced *Sirt1* conditional knockout leukemia-derived cell line under basal conditions or 2-days after 4-Hydroxytamoxifen-induced isogenic loss of *Sirt1*, measured in real time using a Seahorse XF24 instrument. Data are presented as +/- SD of n = 5 wells. **E-F,** Proliferation (left) and quantification (right) of tumor-derived cell lines from ΔE-NOTCH1-induced (E) or HDΔP-NOTCH1-induced (F) *Sirt1* conditional knockout leukemias upon treatment with DBZ or DMSO (control) and ethanol ($Sirt1^{+/+}$) or 4-Hydroxytamoxifen ($Sirt1^{-/-}$) *in vitro*. **G-H,** Quantification (left) and representative flow cytometry plots (right) from of annexin V (apoptotic cells) and 7-AAD (dead cells) staining in tumor-derived cell lines from ΔE-NOTCH1-induced (G) or HDΔP-NOTCH1-induced (H) *Sirt1* conditional knockout leukemias upon treatment with DBZ or DMSO (control) and ethanol ($Sirt1^{+/+}$) or 4-Hydroxytamoxifen ($Sirt1^{-/-}$) *in vitro*. $^{**}P < 0.01$ and $^{***}P < 0.005$ using 1-way analysis of variance (ANOVA).

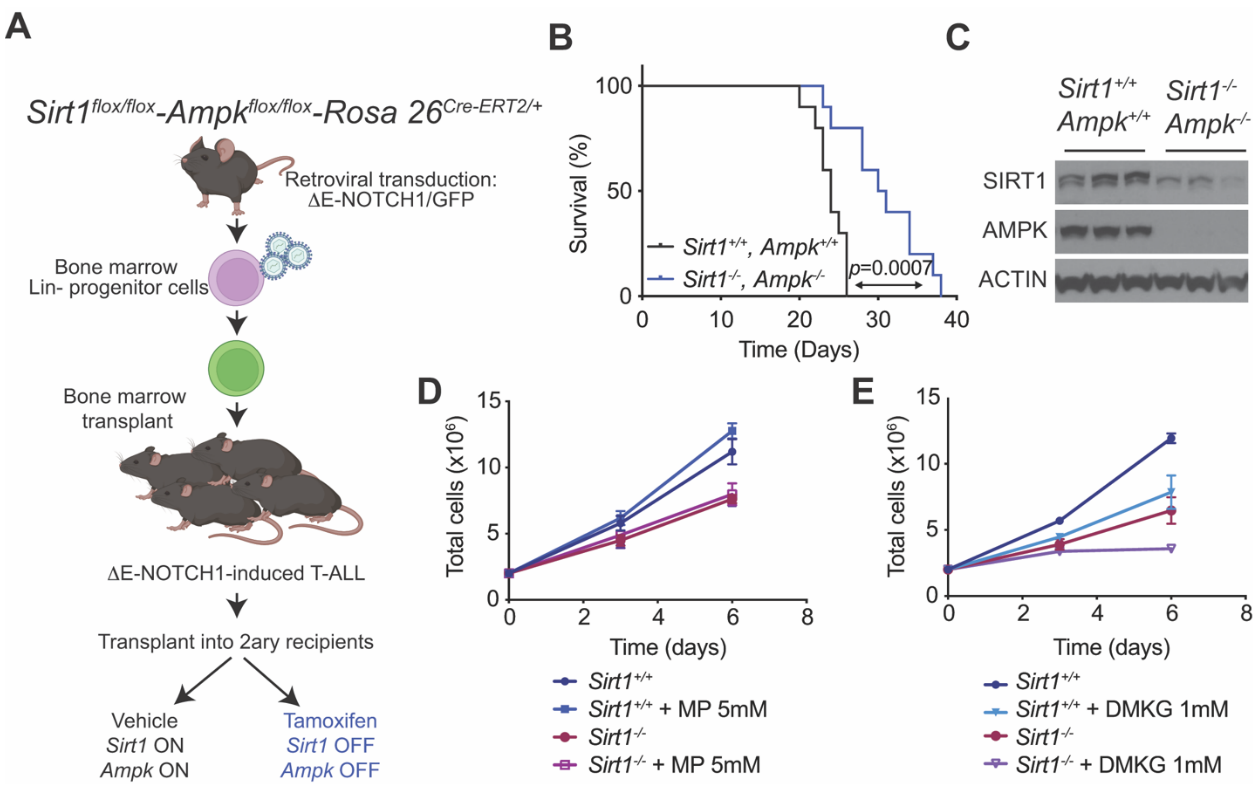


**Figure 4.** Secondary loss of AMPK does not impair the antileukemic effects of SIRT1 loss in T-ALL *in vivo*. **A,** Schematic of retroviral-transduction protocol for the generation of NOTCH1-induced T-ALLs from inducible double *Sirt1/Ampk*-conditional knockout mice, followed by transplant into secondary recipients treated with vehicle or tamoxifen. **B,** Kaplan-Meier survival curves of mice harboring *Sirt1/Ampk*-positive and *Sirt1/Ampk*-deleted isogenic leukemias treated (n = 10 per group; *P* value calculated with log-rank test. **C,** Western blot analysis of SIRT1, AMPK and ACTIN expression in leukemic spleens from terminally ill mice from survival curve in B. **D-E,** Proliferation of a tumor-derived cell line from a ΔE-NOTCH1-induced *Sirt1* conditional knockout leukemia upon treatment with 4-Hydroxytamoxifen ($Sirt1^{-/-}$) or ethanol ($Sirt1^{+/+}$) *in vitro* in the presence of methyl-pyruvate (MP; panel D) or dimethyl-2-oxoglutarate (DMKG; panel E).

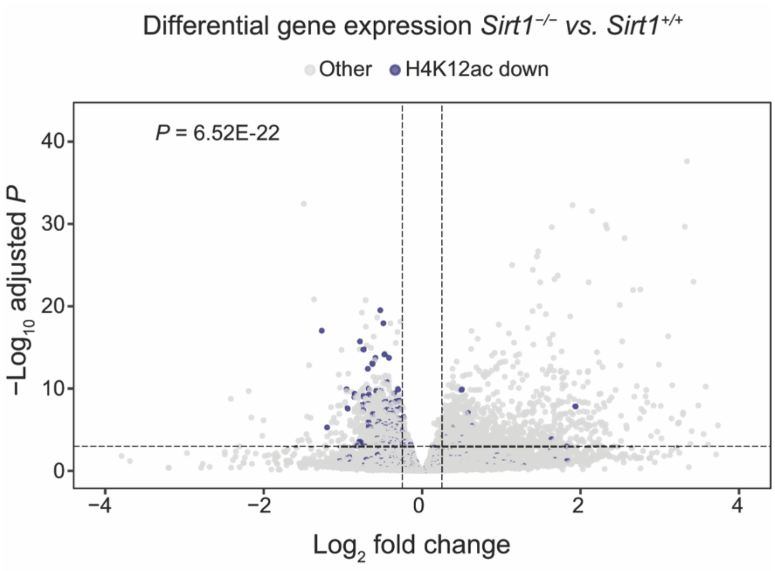


**Figure 5.** Significant correlation between downregulated genes and regions showing reduced H4K12ac upon SIRT1 loss in T-ALL *in vivo*. Volcano plot showing H4K12ac downregulated region genes (shrunken LFC < -0.5) in blue (n=587) and the rest of the regions in light grey. Horizontal dashed line corresponds to the adjusted *P* value threshold of 0.001. Vertical dashed lines correspond to $log_2FC$ changes of +0.25 and -0.25. 102/587 were significantly downregulated at the gene expression level (hypergeometric test *P* value = 6.52E-22)
